# Exploring the K^+^ binding site and its coupling to transport in the neurotransmitter:sodium symporter LeuT

**DOI:** 10.1101/2023.04.04.535525

**Authors:** Solveig G. Schmidt, Andreas Nygaard, Joseph A. Mindell, Claus J. Loland

## Abstract

The neurotransmitter:sodium symporters (NSSs) are secondary active transporters that couple the reuptake of substrate to the symport of one or two sodium ions. One bound Na^+^ (Na1) contributes to the substrate binding, while the other Na^+^ (Na2) is thought to be involved in the conformational transition of the NSS. Two NSS members, the serotonin transporter (SERT) and the *Drosophila* dopamine transporter (dDAT), also couple substrate uptake to the antiport of K^+^ by a largely undefined mechanism. We have previously shown that the bacterial NSS homologue, LeuT, also binds K^+^, and could therefore serve as a model protein for the exploration of K^+^ binding in NSS proteins. Here, we characterize the impact of K^+^ on substrate affinity and transport as well as on LeuT conformational equilibrium states. Both radioligand binding assays and transition metal ion FRET (tmFRET) yielded similar K^+^ affinities for LeuT. K^+^ binding was specific and saturable. LeuT reconstituted into proteoliposomes showed that intra-vesicular K^+^ dose-dependently increased the transport velocity of [^3^H]alanine, whereas extra-vesicular K^+^ had no apparent effect. K^+^-binding induced a LeuT conformation distinct from the Na^+^- and substrate-bound conformation. Conservative mutations of the Na1 site residues affected the binding of Na^+^ and K^+^ to different degrees. The Na1 site mutation N27Q caused a >10-fold decrease in K^+^ affinity but at the same time a ∼3-fold increase in Na^+^ affinity. Together, the results suggest that K^+^-binding to LeuT modulates substrate transport and that the K^+^ affinity and selectivity for LeuT is sensitive to mutations in the Na1 site, pointing toward the Na1 site as a candidate site for facilitating the interaction with K^+^ in some NNSs.

## INTRODUCTION

The family of neurotransmitter:sodium symporters (NSSs) include the transporters responsible for the reuptake of neurotransmitters from the extracellular space following synaptic transmission. Of pronounced interest to neuropharmacology is the subclass of monoamine transporters (MATs), including the dopamine transporter (DAT), the norepinephrine transporter (NET) and the serotonin transporter (SERT), which are molecular targets for a range of psychopharmaceuticals, including drugs against depression, anxiety, neuropathic pain, attention deficit-hyperactivity disorder (ADHD) and narcolepsy^1, 2^. They are also targets for psychostimulant drugs of abuse, such as cocaine and amphetamine^3, 4^. Thus, understanding the molecular basis underlying their ligand binding and transport are of physiological and pharmacological interest.

The NSS member LeuT, a hydrophobic amino acid transporter originating from the thermophile bacterium *Aquifex aeolicus*, was the first NSS structure to be solved^5^. LeuT has served as a structural and mechanistic model for NSS proteins^5, 6^. All NSS proteins are thought to share the LeuT-fold structure, comprising 10 transmembrane (TM) segments ordered in a pseudo-symmetry in the plane of the membrane between the first and the second 5 TMs. The LeuT-fold is mostly followed by two C-terminal TMs. Comparing the available structures stabilized in different states of the transport cycle suggests an overall similar transport mechanism, substantiating LeuT as a valid model protein^7–11^. The substrate binding site is located halfway through the core of the protein and consists of coordinating residues from TM1, 3, 6, 7 and 8^5^. It is flanked by two Na^+^ binding sites, Na1 and Na2. The location and the residues forming the Na1 and Na2 sites are quite conserved between LeuT and the MATs. In the Na1 site, only one of the four coordinating residues differ by a serine to threonine substitution. In the Na2 site, two of the five residues are substituted between LeuT and MATs. A conserved feature shaping the Na^+^ binding sites is the helical unwinding in TM1 and TM6, which exposes backbone-carbonyl oxygen atoms to partake in the ion coordination^5^. In LeuT, the Na1 ion is also coordinated by the carboxyl-group of the substrate, yielding substrate binding highly Na^+^-dependent^5^.

The sodium gradient across the cell membrane is essential for driving substrate uptake in NSS proteins through occupation of the Na^+^ sites. In addition, it has for long been recognized that SERT antiports K^+^ as studies have shown that serotonin uptake was accelerated by an outward directed K^+^ gradient^12^. Consequently, K^+^ antiport was suggested to increase the rate of the return step, which is thought to be the rate-limiting step of the transport cycle^13, 14^. Cryo-EM reconstruction of SERT obtained in KCl revealed an inward-open conformation, suggesting this to be the most prevalent structure with KCl^9^. The resolution of this structure, however, did not allow unambiguous identification of densities for bound ions. Thus, while the conformational details of the K^+^ bound state of SERT are emerging, the location of the K^+^ binding site remains unknown and the antiport mechanism not fully understood. We have shown that K^+^ is antiported also by *Drosophila* DAT (dDAT) by a mechanism that shares similarities with K^+^ antiport in SERT^15^.

Previously, we have reported that LeuT interacts with K^+^ and that the binding favors an outward-closed conformation^16^. Intra-vesicular K^+^ also increased the concentrative capacity of [^3^H]alanine by LeuT, possibly by decreasing substrate efflux, but the details of K^+^’s interaction with LeuT were not fully explored^16^. Here, we use purified LeuT, either in detergent micelles or reconstituted into proteoliposomes, to characterize the molecular components of how K^+^ binds to LeuT, and how it influences the kinetics of substrate transport and the underlying conformational dynamics. Moreover, we mutate Na1 site residues and observe the effect on Na^+^ and K^+^ binding. Analogous to how LeuT for long has served as a structural model for the NSS family of proteins, it here serves as a functional model to suggest a role for K^+^ in the transport mechanism.

## RESULTS

We have previously shown that K^+^ inhibits Na^+^-dependent [^3^H]leucine binding to LeuT and changes its conformational equilibrium^16^. However, fundamental questions remain to be addressed: Is the effect of K^+^ the result of direct binding to a specific binding site in LeuT? If so, where is this site located? What are the kinetic mechanisms underlying the effect of K^+^ on [^3^H]alanine transport? To address this, we expressed His-tagged LeuT in *E. coli*, harvested and solubilized the membranes in n-dodecyl-β-D-maltoside (DDM) and purified the protein by immobilized metal affinity chromatography. Purified LeuT was then used for radioligand binding assays, to analyze the consequences of ion binding on conformational dynamics by transition metal ion FRET (tmFRET), as well as for reconstitution into proteoliposomes for transport assays.

### K^+^ binding is competing with Na^+^ and saturable

To characterize the relationship between Na^+^ and K^+^ binding to LeuT, we investigated the effect of K^+^ on Na^+^-dependent [^3^H]leucine binding using the scintillation proximity assay^17^. Accordingly, we performed a Na^+^ titration with 100 nM [^3^H]leucine (**Figure 1A**). Choline (Ch^+^), seemingly inert to LeuT^16^, was used as counterion to maintain the ionic strength. Na^+^ promoted the binding of [^3^H]leucine to LeuT with an EC_50_ for Na^+^ of 7.7 mM, in line with previous studies^18, 19^. In the presence of K^+^, however, the EC_50_ right-shifted while the B_max_ remained unchanged, consistent with a competitive mechanism of inhibition between Na^+^ and K^+^ as suggested previously^16^.

**Figure 1.**
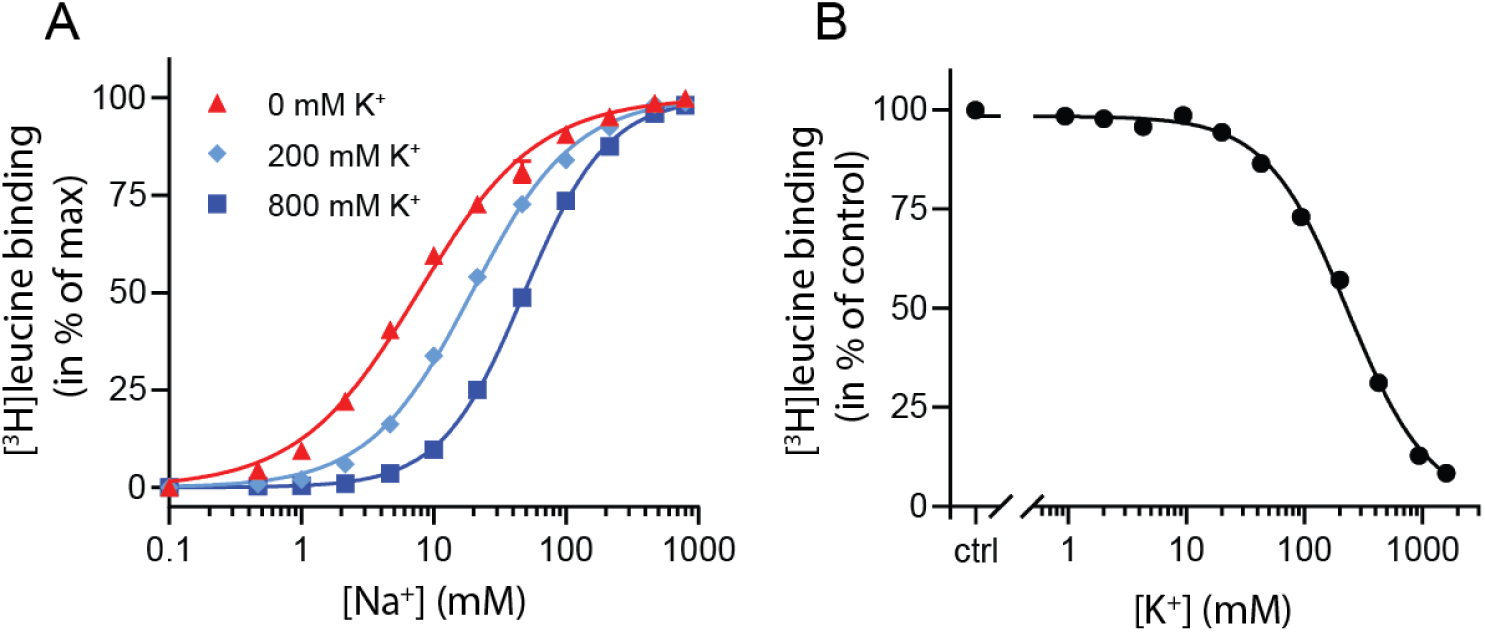
K^+^ competitively inhibits Na^+^-mediated [^3^H]leucine binding in LeuT. (**A**) Na^+^-mediated [^3^H]leucine (10x *K*_d_) binding to purified LeuT in the absence (red triangles) or presence of 200 mM (light blue diamond) or 800 mM (blue squares) K^+^. Data are normalized to B_max_ and fitted to a Hill equation, yielding EC_50_ values in 0, 200 and 800 mM K^+^ of 7.7 [7.3; 8.1], 19.4 [19.1; 19.8] and 48.5 [47.5; 49.5] mM, respectively. (**B**) K^+^-dependent displacement of Na^+^-mediated [^3^H]leucine binding in the presence of 7.7 mM Na^+^, which corresponds to the EC_50_ determined in (A). Data are normalized to a control with 0 mM K^+^ and fitted to a Hill equation with an IC_50_ value of 234.8 [224.8; 243.1] mM. All data points are shown as mean ± standard error of the mean (s.e.m.), n = 3–6 conducted in triplicates. Error bars often smaller than data points. EC_50_ and IC_50_ values are reported as mean [s.e.m. interval]. The ionic strength was maintained by substituting Na^+^ and K^+^ with Ch^+^.

To determine whether the inhibition of [^3^H]leucine binding by K^+^ was saturable, we titrated in K^+^ in the presence of Na^+^ and [^3^H]leucine. Displacement of [^3^H]leucine binding by K^+^ is concentration dependent, with an IC_50_ for K^+^ of 235 [225; 243] mM (mean [s.e.m. interval]) (**Figure 1B**). To ensure that the displacement of [^3^H]leucine by K^+^ was not caused by artefacts originating from the high total salt concentrations (1.6 M), we repeated the displacement assay with a total ionic strength of 208 mM salt using either Ch^+^ or N-Methyl-D-glucamine (NMDG^+^), both of which are inert to LeuT, as counter-ions. Again, K^+^ dose-dependently displaced [^3^H]leucine binding with an inhibition constant not significantly different from that measured in the high salt conditions (**Figure 1-Figure supplement 1**). Although the approach is indirect with respect to K^+^, this saturable, ion-strength independent displacement of [^3^H]leucine binding is indicative of a specific, low affinity K^+^ binding site in LeuT.

### The binding of ions is reflected in changes in the conformational equilibrium of LeuT

To obtain a more direct readout of K^+^ binding to LeuT, we turned to tmFRET. This method relies on the distance-dependent quenching of a cysteine-conjugated fluorophore (FRET donor) by a transition metal (FRET acceptor), here Ni^2+^, coordinated to an engineered α-helical His-X_3_-His site^20^. In LeuT, we have inserted the His-X_3_-His site in extracellular loop (EL)4a (A313H-A317H) and a cysteine at the top of TM10 (K398C) that is labelled with fluorescein-5-maleimide (F5M). The distance between these FRET probes changes upon opening and closing of the extracellular gate in LeuT (**Figure 2A** and **Figure 2 - Figure supplement 1A**). Importantly, this construct, LeuT A313H-A317H-K398F5M (from here and on named LeuT^tmFRET^), retains WT activity with respect to ligand binding affinities^16^. Accordingly, changes in tmFRET intensity is a conformational readout for both ion- and ligand-binding.

**Figure 2.**
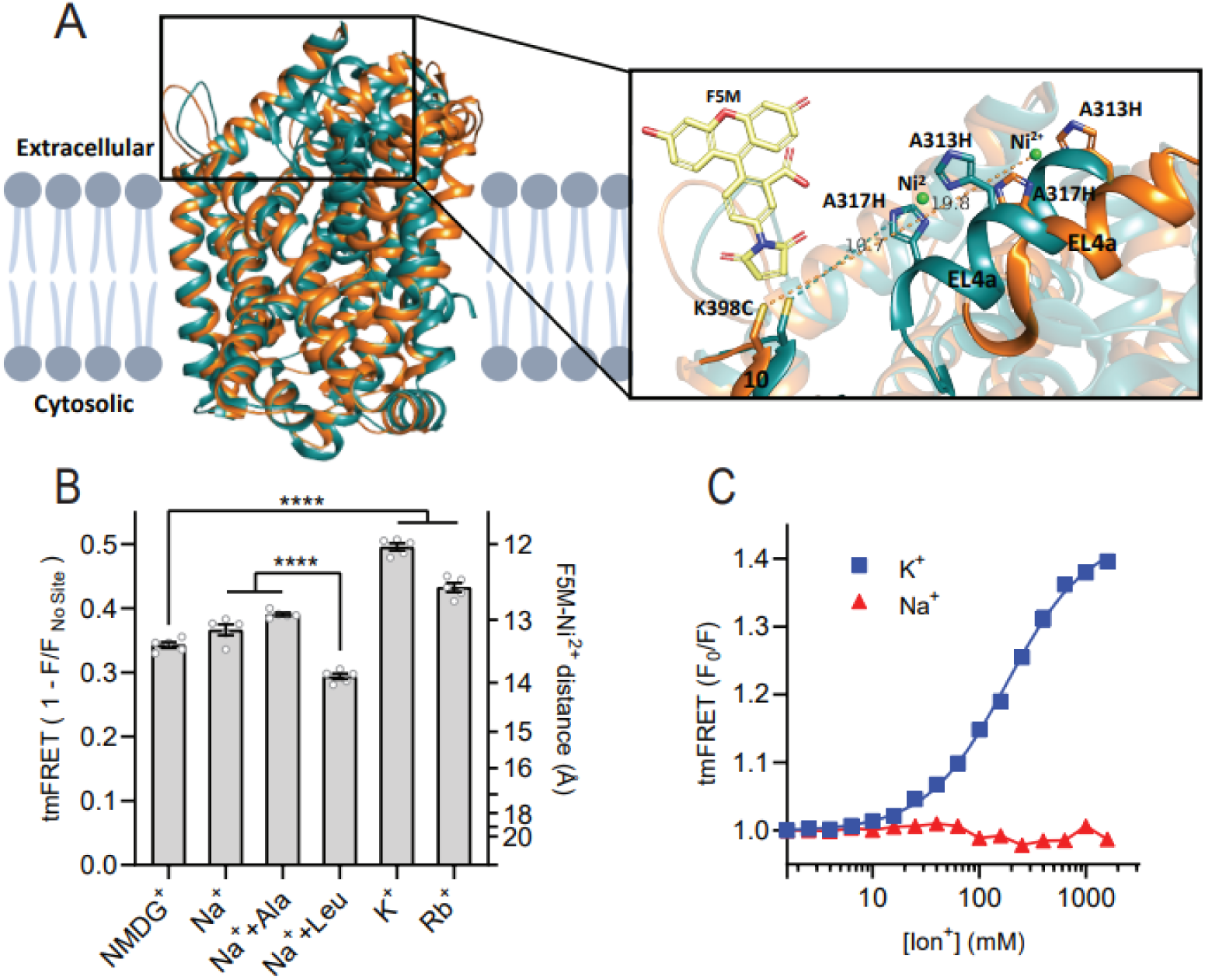
K^+^ shifts the conformational equilibrium towards an outward-closed state. (**A**) Left; cartoon representation of the superimposed structures of LeuT in the outward-open Na^+^ bound state (orange; PDB-ID 3TT1) and inward-open state (turquoise; PDB-ID 3TT3), viewed parallel to the plane of the membrane. Right; enlarged view of the top of TM10 and EL4a, carrying the F5M (yellow, shown adjacent to K398C) and the Ni^2+^-chelating His-X_3_-His site (A313H-A317H). Dashed lines indicate for both structures the distance between the coordinated Ni^2+^ ion (green) and the side chain sulfur of K398C (10.7 Å in 3TT3 and 19.8 Å in 3TT1). (**B**) tmFRET efficiencies (1-F/F_no site_) obtained for LeuT in 800 mM of the ions specified (± 50 µM leucine or 200 µM alanine when indicated) upon saturation of the His-X_3_-His site with 10 mM Ni^2+^. Right vertical axis shows FRET efficiencies converted to distances by the FRET equation. \*\*\*\**p*<0.0001 represents the significance levels from a Tukey multiple comparison-corrected one-way ANOVA. (**C**) tmFRET (F_0_/F) as a function of Na^+^ (red triangles) or K^+^ (blue squares) performed on LeuT^tmFRET^ in the presence of 750 µM Ni^2+^. The K^+^ response fitted to a Hill equation yields a Hill slope of 1.15 ± 0.03 (mean ± s.e.m.) and an EC_50_ of 182.6 [176.4; 189.1] mM, mean [s.e.m. interval]. All data points represent mean ± s.e.m. (error bars often smaller than data points), n = 3-5 conducted in triplicates.

To determine the Ni^2+^ concentrations required to saturate the His-X_3_-His site, we first recorded the FRET efficiency as a function of increasing Ni^2+^ for LeuT incubated with NMDG^+^, K^+^ or Na^+^ ± leucine (**Figure 2 – Figure supplement 1B**). To ensure close to saturating conditions for both Na^+^ and K^+^ while preserving the ionic strength, we applied 800 mM of the ions. The FRET efficiencies at saturating Ni^2+^ concentrations reflect the average distance between the FRET probes and showed that K^+^ uniquely stabilizes a high FRET state, suggesting a shift in the conformational equilibrium of the transporter towards an outward-closed state by K^+^ (**Figure 2 - Figure supplement 1A,B)**. The Ni^2+^ affinity for the His-X_3_-His site is increased approx. 3-fold when Na^+^ or K^+^ is added to LeuT relative to NMDG^+^. This indicates that both Na^+^ and K^+^ bind and stabilize LeuT, including the His-X_3_-His motif, whereas NMDG^+^ does not.

We proceeded by measuring the FRET efficiency only at saturating Ni^2+^ concentrations (10 mM) to obtain FRET efficiencies independent of potential differences in Ni^2+^ affinities. In addition to the conditions above, we measured FRET efficiencies for LeuT incubated in Na^+^ with alanine and in Rb^+^ (**Figure 2B**). Interestingly, applying Rb^+^, a cation often seen to be able to substitute for K^+^ in biological systems, also stabilized a more outward-closed state relative to that in NMDG^+^. The FRET efficiency in the Na^+^/leucine-bound state was decreased relative to that in the Na^+^-bound state, suggesting an, on average, more open-to-out state when adding leucine. We speculate that LeuT adopts a more conformationally restricted equilibrium upon binding of leucine relative to Na^+^ alone, and that this is reflected in a lower FRET efficiency (**Figure 2 - Figure supplement 1A)**. This is also in line with the observation that alanine gives rise to a higher FRET efficiency than leucine (**Figure 2B**) as alanine binding allows a higher degree of conformational freedom in the transporter^18, 21^.

The large difference in tmFRET efficiencies between the apo-state with NMDG^+^ and the K^+^-bound state of LeuT^tmFRET^ allowed us to use the K^+^-induced change in conformational dynamics as a proxy for K^+^ binding. To estimate the affinity for K^+^ to apo-state LeuT, we recorded the FRET efficiencies for LeuT^tmFRET^ incubated with Ni^2+^ in increasing concentrations of K^+^ (**Figure 2C**). We observed an increase in FRET as a function of K^+^ yielding an EC_50_ of 183 [176 ; 189] mM, in line with the affinity determined by K^+^-dependent displacement of Na^+^ and [^3^H]leucine (**Figure 1B**). This EC_50_ is also in line with the affinity from the Schild analysis performed previously^16^. While Na^+^ did not impose major conformational changes, we found that also Rb^+^ induced a conformational response resembling that of K^+^, suggesting that Rb^+^ can indeed substitute for K^+^ although with a lower apparent affinity (**Figure 2 - Figure supplement 1C**). Of note, the high salt concentrations did not affect the intrinsic fluorescence properties of the fluorophore, validating that the responses to titration of the ions were direct results of conformational changes (**Figure 2 - Figure supplement 1D**). Along with the effect of K^+^ on Na^+^-dependent [^3^H]leucine binding, this finding supports the existence of a specific K^+^ binding site in LeuT, and that K^+^ binding to this site induces an outward-closed conformation.

### K^+^ increases the rate of substrate uptake by LeuT

We have previously shown for LeuT reconstituted into liposomes that intra-vesicular K^+^ increases the concentrative capacity of [^3^H]alanine, probably by decreasing its efflux^16^. To expand on these findings and to characterize how substrate transport was affected by the cations in the intra-vesicular buffer, we reconstituted purified LeuT into liposomes containing either Na^+^, NMDG^+^, Cs^+^, Rb^+^ or K^+^ (**Figure 3A**). Under each of these conditions, we measured time resolved [^3^H]alanine uptake driven by a Na^+^ gradient (**Figure 3B**). With equimolar intra- and extra-vesicular Na^+^ concentrations, no [^3^H]alanine transport was observed, indicating that the established Na^+^ gradient did drive [^3^H]alanine uptake (**Figure 3B**). Interestingly, proteoliposomes containing K^+^ displayed a 2.5-fold increase in concentrative capacity compared to those containing Cs^+^ or NMDG^+^ (**Figure 3 – Supplementary table 1**). To ensure that variations in the amount of active LeuT in the proteoliposomes did not affect the uptake capacity, we correlated it to the relative number of active LeuT under each condition (**Figure 3 - Figure supplement 1A**). The concentrative capacity with Rb^+^ was similar to that with K^+^, indicating that Rb^+^ can functionally substitute for K^+^. These data suggest that K^+^, and Rb^+^, are not obligate for LeuT transport, but add a concentrative potential for accumulation of alanine in addition to that obtained solely by the Na^+^ gradient.

**Figure 3.**
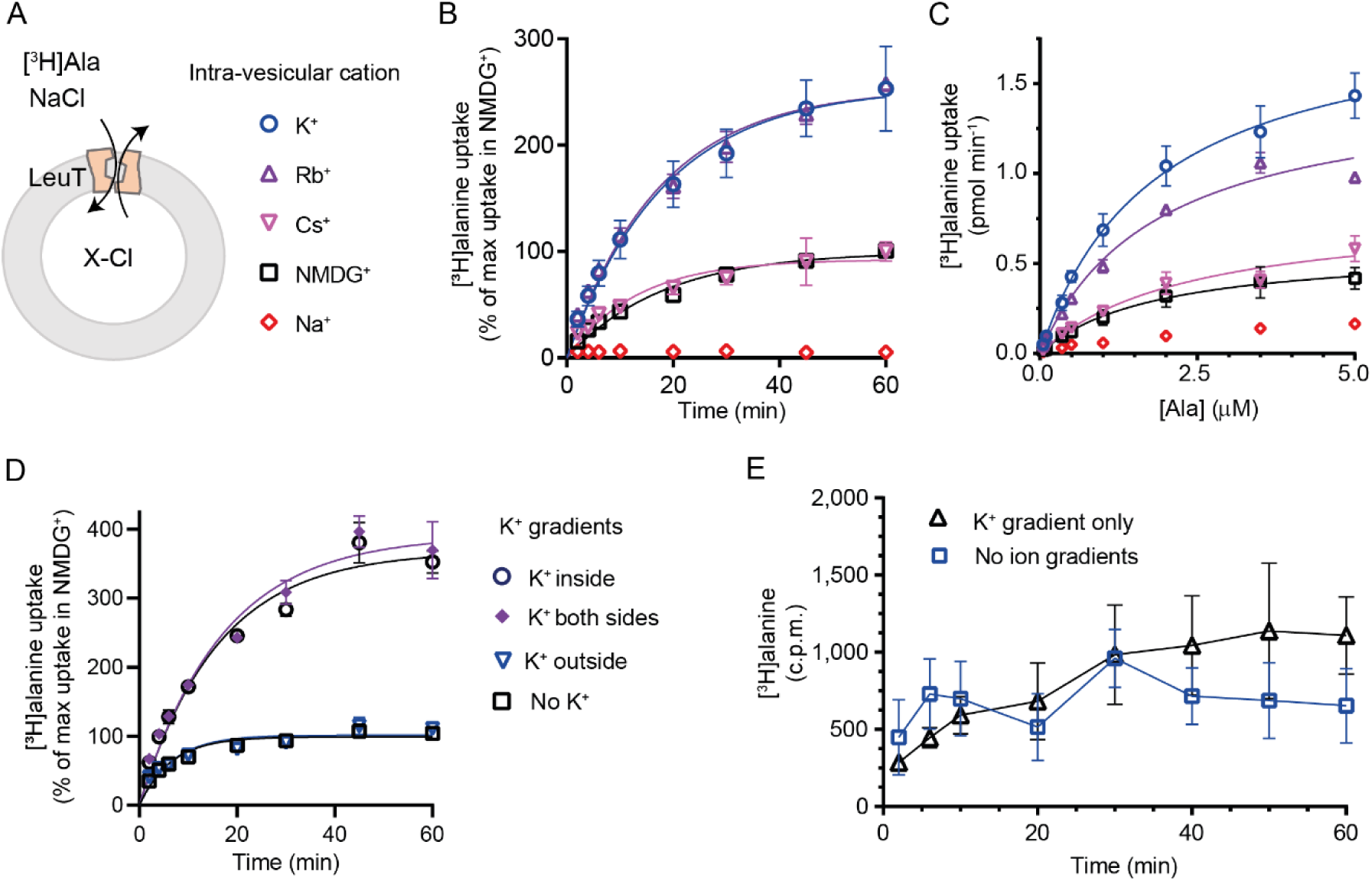
K^+^ modulates LeuT mediated [^3^H]alanine transport. **(A)** Schematic of LeuT reconstituted into liposome containing buffer with various cations (X) and Cl^-^ as corresponding anion. **(B)** Time-dependent [^3^H]alanine (2 µM) uptake in the presence of 200 mM Na^+^ into liposomes containing 200 mM of the various cations (colored as in (A)). Data are fitted to a non-linear regression fit. The maximum uptake predicted with NMDG^+^ is defined as 100% (**supplementary table 1**). **(C)** Concentration-dependent [^3^H]alanine uptake for 5 min (colored as in (A)). Lines are fits to Michaelis-Menten kinetics (**supplementary table 2**). None of the *K*_m_ values were significantly different (Tukey multiple comparison-corrected one-way ANOVA, *p*>0.05). The *V*_max_ value in Cs^+^ was not significantly different from NMDG^+^ (Tukey multiple comparison-corrected repeated-measures one-way ANOVA, *p*=0.5780), but K^+^ and Rb^+^ were significantly different from NMDG^+^ (*p*=0.0011 and 0.0132, respectively). **(D)** Time-dependent [^3^H]alanine uptake in the presence of 50 mM Na^+^ and absence of K^+^ (black squares) was defined as 100%. Addition of 150 mM K^+^ (blue triangles) did not significantly change the maximal uptake (unpaired t-test, *p*=0.92). With 150 mM intra-vesicular K^+^ (blue circles) the maximum uptake increased to 368 ± 15 % (mean ± s.e.m). This increase was not significantly (unpaired t-test, *p*=0.43) affected further when 150 mM K^+^ was also added to the extra-vesicular side (purple diamonds). The ionic strength was kept constant by substitution with NMDG^+^. **(E)** Time-dependent [^3^H]alanine uptake into liposomes containing 25 mM Na^+^ and 200 mM K^+^ in the presence of either 25 mM Na^+^ and 200 mM K^+^ (black line, no ion gradients) or 25 mM Na^+^ and 200 mM NMDG^+^ (blue line, outward directed K^+^ gradient). The data points from the two conditions are not significantly different (unpaired Mann-Whitney test, *p*=0.645). All data points are shown as mean ± s.e.m, n=3 performed in duplicates (B) or triplicates (C-E).

Next, we investigated how the identity of the intra-vesicular cation affected alanine uptake rate. The *K_m_* for [^3^H]alanine in vesicles containing K^+^ was 1.8 ± 0.4 µM, and it was not significantly different upon substitution of intra-vesicular K^+^ with NMDG^+^, Cs^+^ or Rb^+^ (**Figure 3C** and **Figure supplement 3 Table 2**). The *V_max_* for [^3^H]alanine uptake was not significantly different between NMDG^+^-and Cs^+^-containing proteoliposomes. In contrast, the *V*_max_ increased 2.5-3 fold when NMDG^+^ was substituted with Rb^+^ or K^+^. The increased uptake rate could originate from increases in the rates of certain steps in the transport cycle, from a decrease in [^3^H]alanine efflux, or a combination of both.

To gain further insight into the relationship between substrate transport and intra-vesicular K^+^, we investigated how the K^+^ concentration affected *K*_m_ and transport velocity for [^3^H]alanine. Accordingly, LeuT-containing proteoliposomes were prepared in a range of intra-vesicular K^+^ concentrations. We observed that the *V_max_* for [^3^H]alanine transport increased with increasing intra-vesicular K^+^ concentration, whereas the estimated *K_m_* values for alanine were not significantly different (**Figure 3 – Figure supplement 1B** and **Figure supplement 3 table 3**), suggesting that the effect of K^+^ on *V*_max_ increases with increasing occupancy of the K^+^ binding site. To assess how the concentration of extra-vesicular Na^+^ affected the substrate uptake, we measured [^3^H]alanine uptake at initial velocity conditions in increasing concentrations of extra-vesicular Na^+^. We found that the half-saturating extra-vesicular Na^+^ concentration was approximately 25 mM for vesicles containing 200 mM intra-vesicular K^+^, which is in line with the higher affinity for Na^+^ to LeuT compared to K^+^ (**Figure 3 - Figure supplement 1C**).

The increase in [^3^H]alanine transport rate by K^+^ could be due to an imposed driving force by the outward-directed gradient. If so, dissipation of the K^+^ gradient would decrease the [^3^H]alanine transport rate. To allow for the addition of K^+^ on the extracellular side also, we lowered the inward directed Na^+^ gradient to 50 mM. The lowered Na^+^ gradient still resulted in a ∼3.5-fold increased transport capacity with intra-vesicular K^+^ relative to NMDG^+^ (**Figure 3D**). However, dissipation of the K^+^-gradient by application of equal amounts of K^+^ on both sides did not change the transport capacity. Addition of K^+^ only on the outside did also not change the transport capacity relative to that with the Na^+^-gradient only (**Figure 3D**), and only having an outward-directed K^+^-gradient could not drive [^3^H]alanine transport (**Figure 3E**). These results suggest that it is the intra-vesicular K^+^ *per se* that increases the transport rate of alanine and not a K^+^ gradient.

It is difficult to control the directionality of proteins when they are reconstituted into lipid vesicles. They will be inserted in both orientations. Outside-out and inside-out. In the case of LeuT, it is the imposed Na^+^-gradient which determines the directionality of transport. Uptake through the inside-out transporters will probably also happen. Note that the inside-out LeuT will not have the K^+^ binding site exposed to the intra-vesicular environment. Accordingly, a propensity of transporters will likely not be influenced by the added K^+^ and will tend to mask the contribution of K^+^ on the transport mode from the right-side out LeuT. To investigate LeuT directionality in our reconstituted samples, we performed thrombin cleavage of accessible C-terminals on intact and perforated vesicles, respectively. The result suggests that the proportion of LeuT inserted as outside-out is larger than the proportion with an inside-out directionality (**Figure 3 – Figure supplement 1D**).

### Mutations in the Na1 site change the affinity for K^+^

With the elucidation of the impact of K^+^ on LeuT substrate transport, we next sought to identify the binding site for K^+^ in LeuT. Since K^+^- and Na^+^-binding are competitive and K^+^ excludes substrate binding, we chose to focus on the Na1 site (**Figure 4A**). Accordingly, we introduced the following conservative mutations of the amino acid residues in the Na1 site: A22S, A22V, N27Q, T254S and N286Q. The aim was to keep LeuT functional but perturb K^+^ binding. Since it has been shown that H^+^ can substitute for the K^+^ antiport in SERT^22^, we also mutated the adjacent E290, which has been proposed to facilitate H^+^ antiport in LeuT^23, 24–26^. Thus, by substituting it to glutamine (LeuT E290Q), we attempted to mimic the protonated state of E290, which – if the mechanism was similar to that in SERT but facilitated through this not conserved residue – should exclude K^+^ binding.

**Figure 4.**
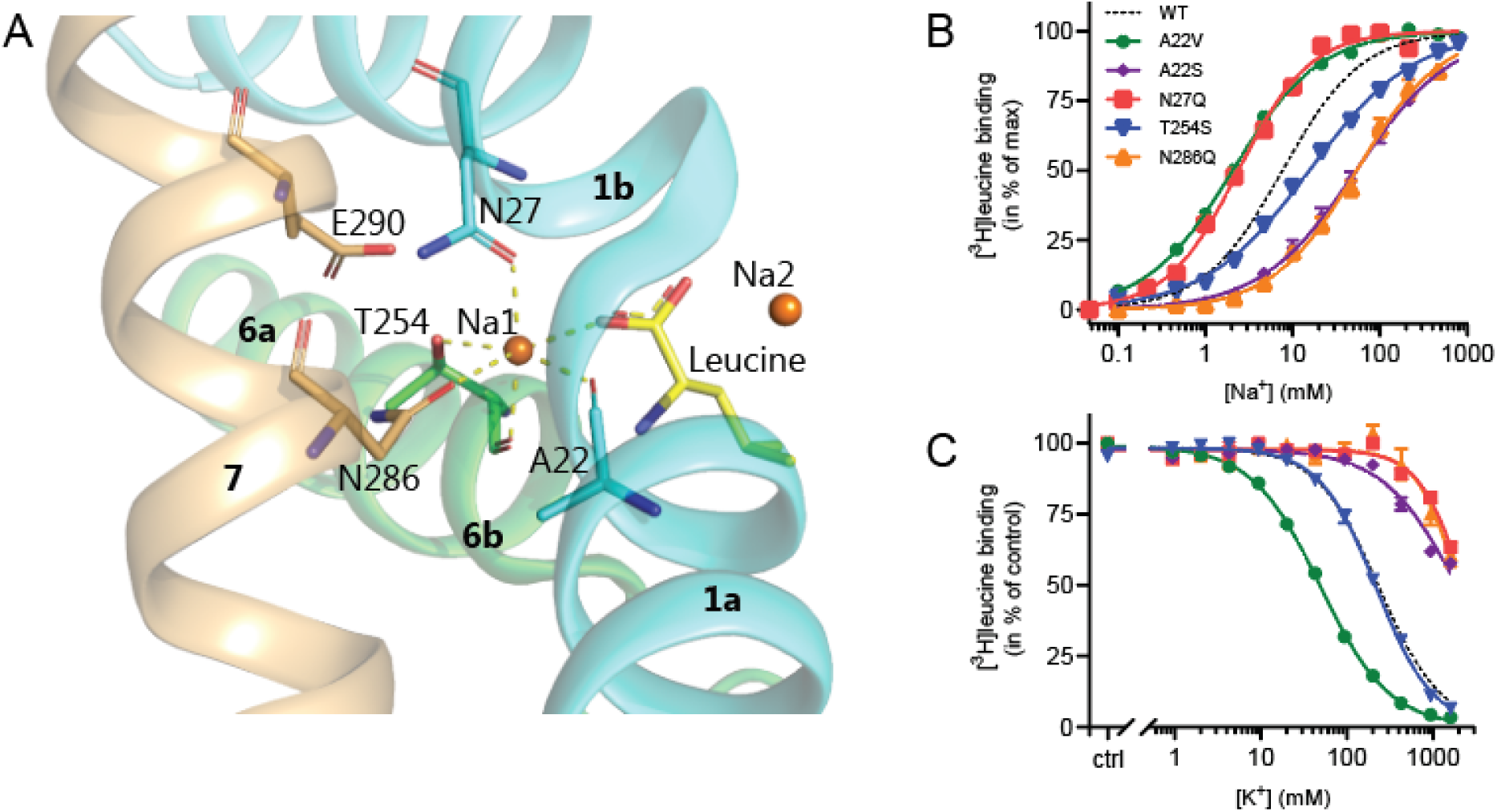
Na1 site mutants of LeuT display altered ion selectivity. (**A**) Cartoon representation of the Na1 site from the outward-occluded LeuT structure (PDB ID: 2A65) with the endogenous Na1 site coordinating residues along with the substrate leucine (yellow) and E290 labeled and shown as sticks. The coordination of Na^+^ is depicted with dashed lines. Structural elements not involved in Na1 site are omitted for clarity. (**B**) Na^+^-mediated [^3^H]leucine (10x *K*_d_) binding for Na1 site mutations, A22V (green circles), A22S (purple rhombi), N27Q (red squares), T254S (blue triangles) and N286Q (orange triangles), with WT shown for reference (black dashed line). Data are normalized to B_max_ and fitted to a Hill equation. (**C**) K^+^-dependent displacement of Na^+^-mediated [^3^H]leucine binding for the Na1 site mutants (colored as in (B)). Na^+^ concentrations equivalent to the EC_50_ values (determined in (B)) and 10x *K*_d_ of [^3^H]leucine for each mutant were used. Data points are normalized to a control without K^+^ and modelled by the Hill equation. All data points are mean ± s.e.m., n = 3-6. The ionic strength was maintained upon substitution with Ch^+^. EC_50_ and IC_50_ values for (B) and (C) are summarized in Table 2.

**Table 2.** Na^+^-mediated [^3^H]leucine binding and the effect of K^+^ addition, and K^+^-dependent Na^+^/[^3^H]leucine displacement in Na1 site mutants. For LeuT Na1 site mutants (and K398C), Na^+^-mediated [^3^H]leucine binding was assayed using 10x *K*_d_ of [^3^H]leucine in the presence of 0, 200, and 800 mM K^+^. For K^+^-dependent displacement of Na^+^-mediated [^3^H]leucine binding, 10x *K*_d_ of [^3^H]leucine and Na^+^ equivalent to the EC_50_ value for each of the mutants were used. Na^+^ and K^+^ were substituted for Ch^+^ to preserve the ionic strength. Data were fitted to a Hill equation, yielding EC_50_ and IC_50_ values reported here as mean [s.e.m. interval], n = 3–6 determined in triplicates. Comparing the Na^+^ EC_50_ obtained for the mutants with that of WT showed significance difference for all (*p*<0.0001) except for E290Q (*p*>0.05), and for the K^+^ IC_50_ all showed significant differences from WT (*p*<0.0001) expect for T254S (*p*>0.05) (Dunnett multiple comparison-corrected one-way ANOVA). N.d. indicates that the value is not determined. WT data are also reported in Figure 1.

| | $EC_{50} Na^+ (mM)$ | $IC_{50} K^+ (mM)$ | $EC_{50} Na^+ + 200 mM K^+ (mM)$ | $EC_{50} Na^+ + 800 mM K^+ (mM)$ | $EC_{50}^{800}/EC_{50}$ |
| --- | --- | --- | --- | --- | --- |
| WT | 7.7 [7.3; 8.1] | 235 [228; 241] | 19.4 [19.1; 19.8] | 48.5 [47.5; 49.5] | ~6 |
| A22V | 2.0 [2.0; 2.1] | 49.0 [47.7; 50.4] | 23.0 [21.7; 24.5] | 70.7 [69.2; 72.2] | ~35 |
| A22S | 53.3 [45.5; 62.5] | > 1900 | 72.4 [62.3; 84.2] | 117 [100; 137] | ~2 |
| N27Q | 2.4 [2.3; 2.5] | > 2300 | n.d. | 4.4 [4.0; 4.8] | ~2 |
| T254S | 15.1 [14.2; 16.0] | 219 [209; 229] | 43.5 [37.4; 50.5] | 90.2 [73.8; 110] | ~6 |
| N286Q | 54.6 [47.5; 62.6] | > 2000 | n.d. | 121 [111; 133] | ~2 |
| E290Q | 9.6 [8.8; 10.4] | n.d. | n.d. | 144 [132; 157] | ~15 |
| K398 | 7.7 [6.8; 8.8] | n.d. | n.d. | 51.5 [48.1; 55.1] | ~7 |

All mutants were expressed in *E. coli* and purified in DDM. They all bound leucine and alanine (**Table 1**). The LeuT mutants A22V, A22S and T254S retained WT-like substrate affinities whereas the mutation E290Q decreased the affinity one order of magnitude and the mutations N27Q and N286Q decreased the affinities about two orders of magnitude. To estimate their Na^+^ affinities, we measured the Na^+^ dependent [^3^H]leucine binding in a next to saturating [^3^H]leucine concentration (10x *K*_d_), thereby taking the differences in substrate affinity for the individual mutants into account (**Figure 4B** and **Table 2**). Mutation of A22V and N27Q increased the Na^+^ affinity by ∼3-fold relative to WT. The T254S mutant caused a 2-fold decrease whereas the A22S and N286Q mutants both decreased the apparent Na^+^ affinity by ∼7-fold. The E290Q mutant retained close to WT Na^+^ affinity (**Table 2**).

**Table 1.** LeuT mutant affinities for leucine and alanine. Affinities were determined by inhibition of [^3^H]leucine with either leucine or alanine. The IC_50_ values, obtained by non-linear regression fit, were converted to *K*_d_ and *K*_i_ values by the Cheng-Prusoff equation. For ^A^-marked *K*_d_’s, affinities were obtained by [^3^H]leucine saturation binding. All experiments were assayed in 200 mM Na^+^ and data are reported as mean [s.e.m. interval].

|  | **Leucine affinity** |  | **Alanine affinity** |  |
| --- | --- | --- | --- | --- |
| | $K_d$ (nM) | $n$ | $K_i$ (μM) | $n$ |
| WT | 6.9 [6.7; 7.2]<sup>A</sup> | 4 | 1.61 [1.58; 1.65] | 4 |
| A22V | 5.4 [4.9; 5.9]<sup>A</sup> | 3 | 1.02 [0.97; 1.08] | 4 |
| A22S | 8.4 [7.7; 9.1]<sup>A</sup> | 3 | 2.11 [2.05; 2.17] | 4 |
| N27Q | 1590 [1530; 1650] | 3 | 255 [248; 263] | 3 |
| T254S | 5.6 [5.4; 5.8]<sup>A</sup> | 3 | 1.17 [1.12; 1.21] | 3 |
| N286Q | 1460 [1390; 1540] | 3 | 287 [277; 297] | 3 |
| E290Q | 193 [169; 221] | 4 | 30.0 [28.9; 31.2] | 3 |

To assess if the mutations in the Na1 site affected the ability to bind K^+^ and test if the competitive mechanism of inhibition was preserved, we repeated the Na^+^ dependent [^3^H]leucine binding experiments in the presence of 800 mM K^+^. For LeuT WT, the added K^+^ caused a ∼6-fold increase in the EC_50_ for Na^+^-dependent [^3^H]leucine binding. Interestingly, mutation E290Q and A22V resulted in increased antagonism by K^+^ relative to WT, causing a 15-fold and 35-fold change, respectively. For the remaining mutants, the antagonism by K^+^ was either less (A22S, N27Q and N286Q) or similar (T254S) (**Table 2** and **Table 2 supplement figure 1**). The observation that the inhibition of Na^+^-dependent ligand binding by K^+^ is retained for LeuT E290Q suggests that binding of K^+^ is not dependent on a negative charge at E290 and can occur in parallel with H^+^ antiport via E290 in LeuT. Also, the fact that the effect of K^+^ on the T254S mutation was indifferent to WT, suggests that the serine residue can completely substitute for threonine in this position.

Next we assessed the impact of the mutations in the Na1 site on K^+^ affinity by measuring the potency by which K^+^ inhibits [^3^H]leucine binding. Again, [^3^H]leucine was added in a near-saturating concentration (10x *K*_d_) and Na^+^ at its determined EC_50_. We observed a marked (∼10-fold) decrease in K^+^ sensitivity for A22S, N27Q, and N286Q, suggesting perturbed K^+^ binding by these mutants (**Figure 4C**). Note that the same N27Q mutant had an increased EC_50_ for Na^+^. In contrast, the potency for K^+^ in inhibiting [^3^H]leucine binding was almost 5-fold increased by the A22V mutation (**Figure 4C** and **Table 2**). The T254S mutation did not alter the K^+^ sensitivity relative to LeuT WT (**Figure 4C** and **Table 2**). The cognate position to Thr254 in SERT is also a serine residue, which could indicate that the threonine to serine substitution is tolerated in terms of K^+^ sensitivity between NSSs from different species.

Taken together, the Na1 site mutations showed a marked and differentiated response to the effects of both Na^+^ and K^+^. For some mutants, the affinities for both Na^+^ and K^+^ were decreased (A22S and N286Q) or increased (A22V). For one, the effects were differentiated so that the Na^+^ affinity was increased, while the K^+^ affinity was decreased (N27Q).

### The conformational responsiveness is altered in the Na1 site mutants

With coordinates from TM1, 6 and the bound substrate, the Na1 site is likely a central mediator of conformational changes. To investigate how the conformational equilibria in the transporters were affected by the Na1 site mutations, we introduced them into the LeuT^tmFRET^ background and probed their response to Na^+^, K^+^ and substrate binding with respect to changes in tmFRET. We first investigated the conformational equilibria in N286Q^tmFRET^ in NMDG^+^ (apo state), in Na^+^ with and without leucine, and in K^+^. Surprisingly, even though the EC_50_ and IC_50_ values for Na^+^ and K^+^, respectively, were markedly increased (decreased affinities), the conformational equilibria of N286Q resembled that of LeuT WT (**Figure 5A**). This suggests that the substantial decreases in ion affinities and selectivity, imposed by the asparagine to glutamine substitution, do not result from conformational biases, but likely from a direct mutual modulation of their binding site. However, we failed to observe any [^3^H]alanine transport activity by reconstituted LeuT N286Q, likely as a result of its low substrate affinity.

**Figure 5.**
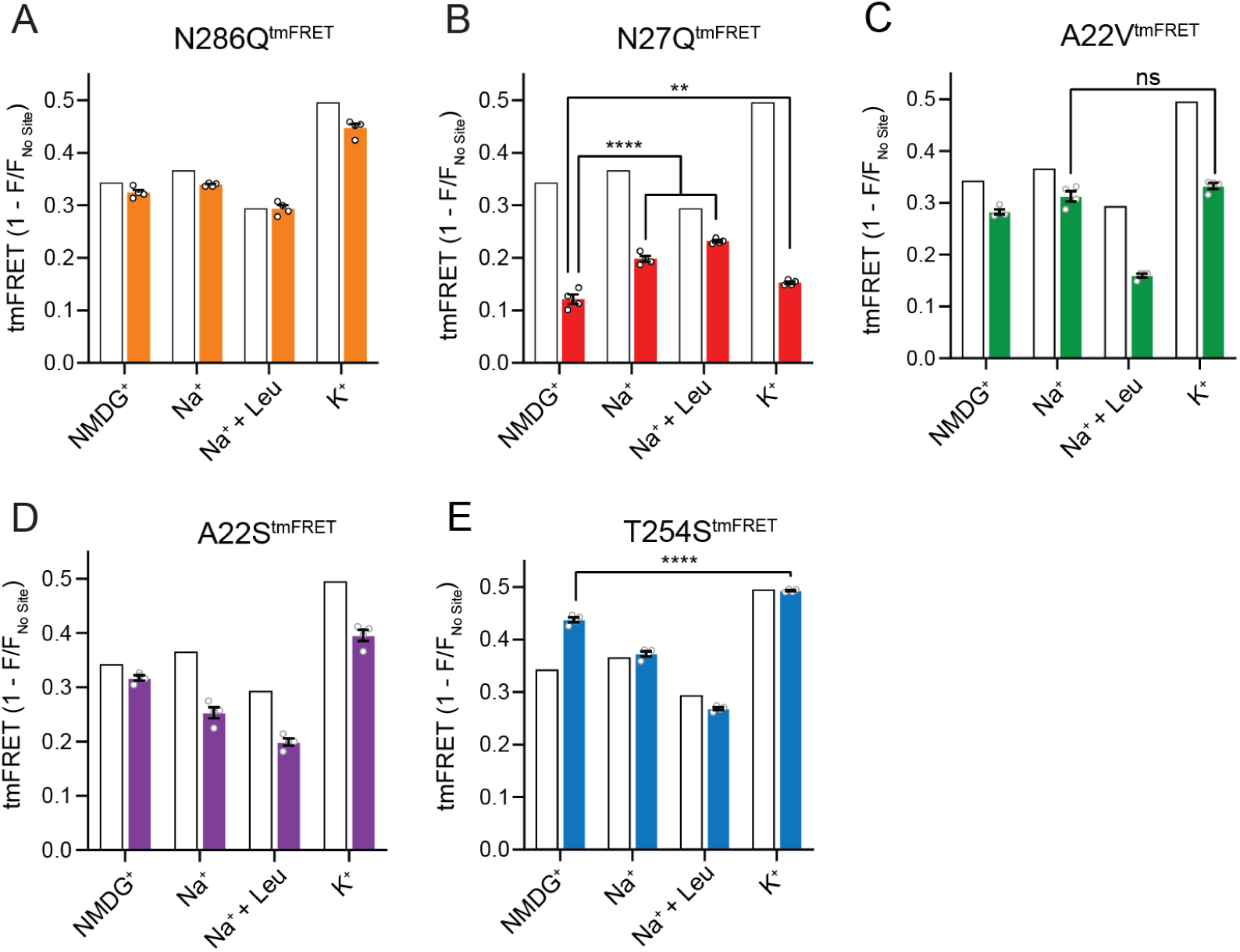
The Na1 site LeuT mutants exhibit different conformational equilibria. (**A-E**) tmFRET efficiencies (1-F/F_no site_) for N286Q^tmFRET^ (orange, **A**), N27Q^tmFRET^ (red, **B**), A22V^tmFRET^ (green, **C**), A22S^tmFRET^(purple, **D**), and T254S^tmFRET^ (blue, **E**) incubated in 800 mM of the indicated ions and 50 µM for leucine. The His-X_3_-His site was saturated with 10 mM Ni^2+^. The corresponding tmFRET efficiencies for LeuT^tmFRET^ (white bars) are shown for reference. All data points are mean ± s.e.m., n = 4, performed in triplicates. In (B), \*\**p*<0.001; \*\*\*\**p*<0.0001 represent the significance levels from a Tukey multiple comparison-corrected one-way ANOVA, comparing the mean obtained in K^+^ with that in NMDG^+^, and the mean in Na^+^ and Na^+^ ± leucine with that in NMDG^+^. In (C) and (E), ns, not significant and \*\*\*\**p*<0.0001 using a one-way ANOVA with Bonferroni multiple comparison correction.

The N27Q^tmFRET^ showed a markedly lower tmFRET efficiency in the apo state compared to WT (**Figure 5B**), suggesting that the mutation biases the conformational equilibrium towards a more outward-open conformation. Addition of Na^+^ and leucine restored the conformational equilibrium towards significantly more outward-closed states, but not to the same extent as found for the LeuT^tmFRET^ construct. However, we hardly observed any change in the tmFRET efficiency with K^+^ in the N27Q mutant. This is in line with its marked decrease in K^+^ affinity. The result could either suggest that Asn27 is important for the binding of K^+^, or that the mutation causes a conformational bias which makes this mutant rarely visit the K^+^-selective state. We were unable to observe any [^3^H]alanine transport activity by the LeuT N27Q mutant when reconstituted into K^+^-containing vesicles.

Mutant A22V^tmFRET^ displayed WT-like tmFRET efficiency in NMDG^+^ and Na^+^, but the tmFRET efficiency induced by K^+^ was not significantly different from that induced by Na^+^ (**Figure 5C**). This indicates that the conformational equilibrium of A22V^tmFRET^ apo-form is not changed by the mutation. In addition, although both Na^+^ and K^+^ ions bind with a higher affinity to LeuT A22V than to WT (**Figure 4**), the tmFRET data suggests that K^+^ no longer promotes the conformational shift towards the outward-closed conformation. To further evaluate this, we reconstituted the mutant into liposomes and measured its ability to transport alanine (**Figure 5 – Figure supplement 1A**). With equimolar Na^+^ on each side of the lipid bilayer, we observed a specific [^3^H]alanine signal, which could either reflect transport driven by the [^3^H]alanine gradient alone or simply binding of [^3^H]alanine to LeuT. In the presence of a Na^+^ gradient, we observed a minor, but significant increase in specific [^3^H]alanine counts. However, the substitution to intra-vesicular K^+^ did not affect the [^3^H]alanine activity. Further investigations must clarify whether the changes in observed [^3^H]alanine activity constitute a transport- or a binding event.

The LeuT A22S mutant displayed a decrease in both Na^+^ and K^+^ affinity (**Figure 4**). When inserting it into the LeuT^tmFRET^ background (A22S^tmFRET^), the pattern in tmFRET efficiencies were largely unaltered from LeuT^tmFRET^, although with a minor reduction in responses upon addition of ligands (**Figure 5D**). This suggests that the mutation only has minor effect on the overall conformational equilibrium independent of the added ligands. When inserted into liposomes, LeuT A22S retained the ability to transport alanine in the presence of a Na^+^ gradient. As for LeuT WT, intra-vesicular K^+^ increased the concentrative capacity for [^3^H]alanine transport. However, the increase was 3-fold higher than what we observed for LeuT WT (**Figure 5 – Figure supplement 1B**). The WT-like conformational equilibrium could suggest that the decreased Na^+^ and K^+^ affinities are due to a direct perturbation of the ion binding site by the A22S mutation.

Finally, we characterized the tmFRET response for T254S^tmFRET^. The mutant exhibited a slightly higher tmFRET efficiency in the apo state (NMDG^+^), but its conformational response to Na^+^, substrate and K^+^ binding was WT-like (**Figure 5E**). Reconstituted into liposomes, LeuT T254S transported [^3^H]alanine and retained a WT-like increase in concentrative capacity with intra-vesicular K^+^ (**Figure 5 - Figure supplement 1C**). As the substitution is the only apparent difference between the Na1 site in LeuT relative to the human SERT, DAT and NET it could suggest a similar functionality of the Na1 site by the four transporters.

In all, although we are unable to discern between direct and indirect effects imposed by the mutants, our results do both reflect concerted and opposed consequences on Na^+^ and K^+^ binding, conformation, and substrate transport.

## DISCUSSION

In this study, we have examined the binding of K^+^ ions to purified LeuT stabilized in detergent micelles. We have determined the binding potency through competition binding with Na^+^ and radiolabeled ligand and by changes in the conformational equilibrium of the transporter induced directly by K^+^ as well as explored the role of K^+^ in the transport process with LeuT reconstituted into liposomes. Additionally, we have investigated the binding site for K^+^ by a mutational screen of the residues contributing to the already known sodium binding site, Na1, which is conserved among NSSs.

To define an interaction between a ligand and a protein as being the result of a binding site, it must be saturable. Here, we show that K^+^ binding is saturable both when assessing its inhibition of Na^+^-dependent [^3^H]leucine binding and when applying tmFRET as a direct conformational readout. The affinity is around 180 mM measured with tmFRET and by the Cheng-Prusoff equation, applied for competitive inhibitors, the IC_50_ from the Na^+^-dependent [^3^H]leucine binding converts to a *K*_i_ of 124 [122;127] mM. This is in the same range as reported previously^16^. Even though ions are quite abundant in many biological systems, an affinity above 100 mM is rather low. We can only speculate whether this is within a physiological range for *A. aeolicus*. However, even at low occupancy, eg. at its *K*_i_ when occupancy is 50 %, the bound K^+^ would regulate the transport velocity.

TmFRET provides a means for a direct measurement of K^+^ binding to LeuT based on changes in the conformational equilibrium of the ensemble of transporters. The tmFRET efficiency in this study reflects intramolecular distance changes between the extracellular side of TM10 and EL4. According to the solved LeuT crystal structures, it predicts that this distance will gradually decline upon transition from the outward open, through the occluded, to the inward open conformation^5, 7^. K^+^ specifically induces a high FRET efficiency relative to those of the other ions and ligands tested, and titration with K^+^ fits a Hill model with a slope of 1.15 ± 0.03 (mean ± s.e.m.), which is in accordance with the existence of one K^+^ binding site in LeuT that - when occupied - biases the conformational equilibrium to an outward-closed state. Interestingly, Rb^+^ induced a response similar to that of K^+^, which correlates with previous observations that Rb^+^ can substitute for K^+^ binding due to similarities in size and preferred coordination^16^. The decreased FRET efficiency in the Na^+^/leucine-bound state, relative to that in the Na^+^-bound state, could suggest that the addition of leucine on average favors a more open-to-out state, although this contradicts with the known crystal structures. However, we speculate that instead this result could be a consequence of the principle by which steady-state FRET efficiencies for heterogeneous dynamic ensembles are biased towards shorter distances^20, 27, 28^. If LeuT adopts a more conformationally restricted equilibrium upon binding of Na^+^ and leucine relative to Na^+^ alone, thereby lowering the frequency by which the transporter visits the outward-closed state, this could reflect the lower FRET efficiency observed for the Na^+^/leucine bound state.

When we reconstituted LeuT into liposomes, intra-vesicular K^+^ increased the capacity and *V*_max_ for [^3^H]alanine uptake compared to intra-vesicular NMDG^+^ and Cs^+^. Rb^+^ displayed a similar effect, suggesting that the conformational effect of Rb^+^ measured by tmFRET translates to an effect on transport function. Dissipating the potassium gradient or adding K^+^ solely on the extra-vesicular side of the proteoliposomes, did not affect uptake capacity. Neither were a potassium gradient alone able to drive uptake of [^3^H]alanine. This suggests that the effect of K^+^ on LeuT uptake is independent of a K^+^ gradient. Conservative mutations in the Na1 site and the adjacent residue Glu290 retained the ability of the transporter to bind ligands, but only A22S and T254S showed sustained [^3^H]alanine transport into proteoliposomes, and a persistent functional effect of K^+^. Interestingly, the LeuT N27Q possessed no apparent affinity for K^+^ but increased Na^+^-affinity. Also, the tmFRET data suggested it irresponsive to K^+^, but also showed an altered conformational equilibrium. This could suggest that the N27Q mutation causes a bias towards an outward-open conformation. [^3^H]Alanine transport by LeuT A22V showed no effect by the addition of intra-vesicular K^+^. This correlates with the loss of apparent K^+^-induced conformational response as assessed in the tmFRET studies (**Figure 5**). However, even though the mutant showed Na^+^-dependent [^3^H]alanine transport, the rate constant was high (1.55 min^-1^) relative to LeuT WT (0.056 min^-1^). This could suggest that the A22V mutation influences additional functional properties, such as entering an exchange mode. Further investigations are needed to clarify this.

Despite the known role of K^+^ in the transport mechanism of SERT and other NSSs, only few studies have attempted to localize the K^+^ binding site^16, 29, 30^. Mutations of Asn338 (corresponding to Asn286 in LeuT) in the Na^+^- and K^+^-coupled amino acid transporter KAAT1 from *M. sexta* suggest that this residue is important for cation selectivity and coupling^29^. Molecular dynamics simulations of LeuT showed that the K^+^-ion could jump between the Na2 and Na1 sites^30^. Mutation in the Na2 site of LeuT (T354V) resulted in a transporter locked in the outward-closed state, and mutation in the Na1 site (T254V) resulted in a transporter which showed a decreased conformational response to K^+16^.

To the best of our knowledge, this work represents the first systematical examination of K^+^ binding in the Na1 site. We found mutations that either increase or decrease the affinities for both Na^+^ and K^+^, but we also saw mutations with opposite effects with respect to the affinities, and thereby selectivity, for the two ions. These changes in ion affinities were independent of conformational biases as assessed with tmFRET. This decoupling in ion selectivity and conformational bias suggests that the alterations result from direct modulations of the binding site for K^+^. However, binding of the two Na^+^ ions and substrate in LeuT is synergistic, making it difficult to completely isolate effects of mutations to a specific site. Consequently, we cannot exclude the possibility that K^+^ binding could also involve the Na2 site or another unknown site. Nevertheless, we consider our mapping of the effect on K^+^ binding by mutations in Na1 a robust starting point for the quest to identify the K^+^ binding site in NSSs.

We propose that K^+^ binding either facilitates LeuT transition from inward- to outward-facing (the rate limiting step of the transport cycle), or solely prevents the rebinding and possible efflux of Na^+^ and substrate. It could also be a combination of both. Either way, intracellular K^+^ will lead to an increase in V_max_ and concentrative capacity. Note that our previous experiment showed an increased [^3^H]alanine efflux when LeuT transports alanine in the absence of intra-vesicular K^+16^. Specifically, the mechanistic impact of K^+^ could be to catalyze LeuT away from the state that allows the rebinding of Na^+^ and substrate. This way, K^+^ binding would decrease the possible rebinding of intracellularly released Na^+^ and substrate, thereby rectifying the transport process and increase the concentrative capacity and *V*_max_ (**Figure 6**). Our results suggest that K^+^ is not counter-transported but rather promotes LeuT to overcome an internal rate limiting energy barrier. However, further investigations must be performed before any conclusive statement can be made here. If K^+^ is not counter-transported, LeuT might comply with the mechanism previously suggested for the human DAT^31^.

**Figure 6.**
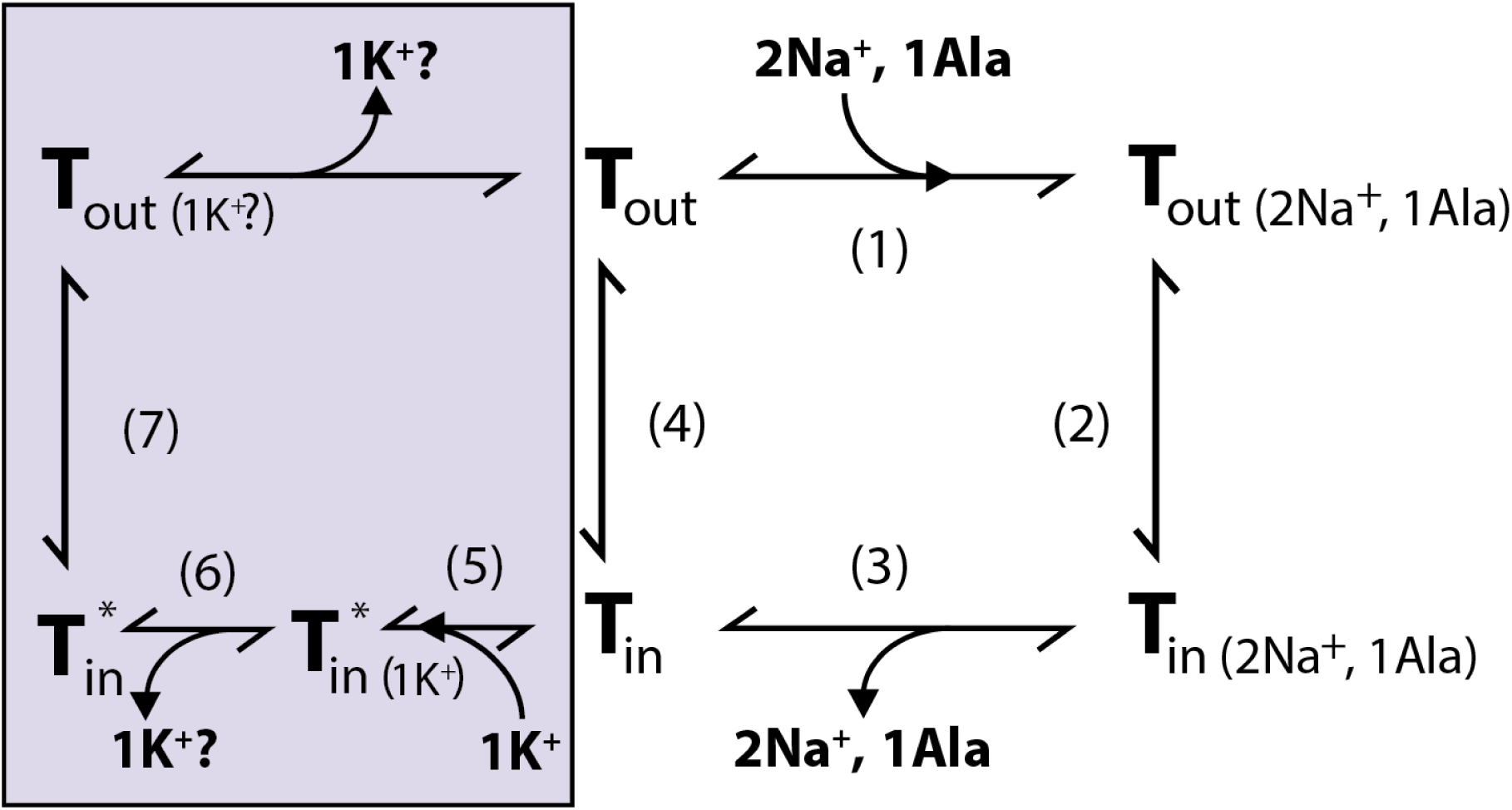
Proposed role of K^+^ in the translocation cycle. Here exemplified for alanine (Ala) as substrate. LeuT apo-form will likely reside in an equilibrium between its outward- (**T**_out_) and inward-facing (**T**_in_) conformation. After Na^+^ and Ala are bound (1) and released (3), LeuT can either i) rebind Na^+^ and Ala, which will promote efflux (2); ii) transition to the outward-open conformation in its apo-form (4), or iii) bind K^+^ (5). K^+^ binding will promote an inward-facing LeuT state (T*) which is unable to bind Na^+^, either by competitive inhibition or by promoting a state that does not allow Na^+^ binding. From here, K^+^ could either be released again to the intracellular environment (6) or be counter-transported (7).

Taken together, K^+^ binding seems conserved between LeuT, SERT^32, 33^, dDAT^15^, and potentially human DAT^15, 34^. Also, the ability to induce a conformational change toward an outward-closed/inward open state appears to be a common mechanism^9, 15, 33, 35^. The conservation of K^+^ binding, but lack of K^+^ antiport, has also been observed in a bacterial member of the SLC1 carrier family^36, 37^. Perhaps regulation of Na^+^-dependent substrate transport by K^+^ is a more common mechanism than previously anticipated.

## MATERIALS AND METHODS

### Reagents

Unless otherwise stated, reagents were purchased from Merck (Life Sciences).

### Cloning of LeuT Mutants

DNA encoding LeuT from *Aquifex aeolicus,* fused C-terminally to a thrombin protease cleavage site and an octahistidine tag, was cloned into a pET16b vector. Single mutations in the Na1 site (A22V, A22S, N27Q, T254S, N286Q, E290Q) were inserted into LeuT constructs with pre-existing pairs of mutations (A313H-A317H-K398C; K398C) and without (WT). Full gene sequences were verified by DNA sequencing (Eurofins Genomics).

### Expression and purification of LeuT

Expression and purification of LeuT variants containing the tmFRET pair mutations were performed essentially as described previously^16^. In brief, *E. coli* C41(DE3) cells were transformed with pET16b plasmid encoding the desired LeuT Na1 site variants and single colonies were cultivated at 37°C in Lysogeny Broth. Expression was induced at OD_600_ ῀ 0.6 upon induction with β-D-1-thiogalactopyranoside (IPTG). Harvested cells were disrupted using a cell disruptor (CF1, ConstantSystems) and isolated crude membranes were solubilized with 1.5 % (w/v) n-dodecyl-β-D-maltopyranoside (DDM) (anagrade, Anatrace). Solubilized LeuT was incubated with Ni^2+^-NTA resin (Thermo Fischer Scientific) and 40 mM imidazole, batch-washed and labelled overnight with fluorescein-5-maleimide (F5M) (Thermo Fischer Scientific). Following wash of the resin with 15 successive column volumes of buffer (20 mM Tris-HCl (pH 7.4), 20 % (v/v) glycerol, 200 mM KCl, 0.05 % (w/v) DDM, 100 µM tris(2-carboxyethyl)phosphine (TCEP) (hydrochloride solution) containing 90 mM imidazole, immobilized LeuT was eluted and frozen in buffer with 340 mM imidazole. Labeling efficiency and specificity were examined by absorbance measurements (280 and 490 nm) and SDS-PAGE analysis, respectively. For LeuT variants on WT background (devoid of tmFRET mutations), the F5M labelling step was skipped and the resin washing procedure was performed with 3 and 5 column volumes of buffer containing 60- and 90 mM imidazole, respectively. For reconstitution in liposomes, LeuT was dialyzed overnight in buffer (20 mM Tris-HCl (pH 7.4), 20 % (v/v) glycerol, 200 mM KCl, 0.05 % (w/v) DDM) to remove imidazole. The protein was subsequently concentrated to > 2 mg/ml.

### Pharmacological characterization of LeuT mutants

Pharmacological studies were conducted on purified LeuT variants by virtue of the Scintillation Proximity Assay (SPA). Leucine affinities for WT LeuT and mutants displaying WT substrate affinities (A22V, A22S) were determined by saturation binding of [^3^H]leucine (25 Ci/mmol) (PerkinElmer), with unspecific binding corrected for by the addition of 100 µM unlabelled leucine. Alanine and leucine affinities for the remaining LeuT variants were determined by the ability of increasing concentrations of un-labelled alanine or leucine, respectively, to competitively displace a fixed concentration of [^3^H]leucine. Assayed in a 96-well plate (Corning), LeuT was mixed with Yttrium Silicate Copper (YSi-Cu) His-tag SPA beads (PerkinElmer) and [^3^H]leucine in binding buffer (200 mM NaCl, 20 mM Tris (pH 8), 20 % (v/v) glycerol, 0.05 % (w/v) DDM and 100 µM TCEP). The following conditions were applied for the mutants tested: WT, K398C, A22V, A22S: 0.3 µg ml^-1^ protein, 1.25 mg ml^-1^ YSi-Cu His-tag SPA beads, 120 nM [^3^H]leucine (25 Ci mmol^-1^); N27Q, N286Q: 3 µg ml^-1^ protein, 1.6 mg ml^-1^ YSi-Cu His-tag SPA beads, 1200 nM [^3^H]leucine (4 Ci mmol^-1^); E290Q: 1.5 µg ml^-1^ protein, 1.6 mg ml^-1^ YSi-Cu His-tag SPA beads, 200 nM [3H]leucine (20 Ci mmol^-1^). The Na^+^ dependency on [^3^H]leucine binding was determined by mixing LeuT, YSi-Cu His-tag SPA beads (in equivalent concentrations as above) and a fixed concentration of [^3^H]leucine (10 x *K*_d_, as determined in 200 mM NaCl) in binding buffer supplemented with increasing concentrations of Na^+^. The specific activity of [^3^H]leucine was kept approximately inversely proportional to the concentration of [^3^H]leucine and protein used. This experiment was repeated in the presence of 200 and 800 mM K^+^. The K^+^ - dependent inhibition of Na^+^ -mediated [^3^H]leucine binding was assayed by subjecting LeuT mutants (double the concentration as above) to increasing concentrations of K^+^ (0-1600 mM) in the presence of Na^+^ equivalent to the EC_50_ determined with 10 x *K*_d_ of [^3^H]leucine. The unspecific binding of [^3^H]leucine was determined upon addition of 100 µM (WT, A22V, A22S) or 300 µM (N27Q, N286Q) of unlabelled leucine. Ionic strengths were preserved by substituting Na^+^ and K^+^ for Ch^+^. For competition binding and ion dependent [^3^H]leucine binding experiments, ligand depletion was avoided by maintaining ≥20-fold molar excess of [^3^H]leucine relative to LeuT. Sealed plates were incubated for ∼16 hours at 4°C with gentle agitation and counts per minute (c.p.m.) were recorded on a 2450 MicroBeta^2^ microplate counter (PerkinElmer) in “SPA” mode.

### Analysis of SPA Data

Saturation binding data were corrected for unspecific binding, normalized to *B*_max_ and fitted to a one-site binding regression from which *K*_d_ values were obtained using GraphPad Prism 9.0. For competition binding, data points were normalized to a control (without competing unlabelled leucine or alanine) and fitted to a single-site log(inhibitor)-response model. Derived IC_50_ values (inhibitor concentration that reduces binding of radioligand by 50 %) were converted to inhibition constants *K*_i_ by the Cheng-Prusoff equiation:

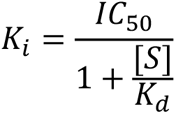

Where [S] and *K*_d_ refer to the concentration and affinity, respectively, of the radioligand. Data points for K^+^ competition binding were corrected for unspecific binding and normalized to a control (absent for K^+^), whereas data for Na^+^-dependent [^3^H]leucine binding were normalized to the maximum response predicted by the model. Both were fitted to the Hill equation from which IC_50_ and EC_50_ values, respectively, were extracted. All experiments were performed at least three independent times in triplicates for specific and triplicates/duplicates for unspecific binding. Data points and Hill slopes are reported as mean ± s.e.m. and *K*_d_, *K*_i_ and EC_50_ values are reported as [s.e.m. interval]. Statistical analysis was performed with a *post hoc* Dunnett multiple test in a one-way analysis of variance, comparing every mean with that obtained in the absence of competitor.

### Transition Metal ion FRET

LeuT Na1 site mutants examined with tmFRET were encoded with a set of FRET pair mutations (A313H-A317H-K398C / K398C) designed and characterized previously^16^. Purified and fluorescein-labelled LeuT variants were centrifuged at 15,000 *g* at 4°C for 10 min, and diluted to ∼10 nM (adjusted for labelling efficiency) in fluorescence buffer (20 mM Tris-Cl (pH 8), 0.05 % (w/v) DDM, 100 µM TCEP), supplemented with 800 mM chloride salts of NMDG^+^, Na^+^ ± 50 µM leucine/ 200 µM alanine, K^+^ or Rb^+^ as specified. Samples were incubated for 30 min at room temperature in the dark. If not otherwise specified, parallel experiments on LeuT constructs with and without the His-X_3_-His motif were conducted, here referred to with the suffixes ^tmFRET^ and ^K398C^, respectively. Fluorescence measurements using a single saturating concentration of Ni^2+^ were assayed in a Swartz cuvette (Hellma Analytics) inserted into a FluoroMax-4 spectrophotometer (HORIBA Scientific) temperature controlled to 25°C. Reference-corrected fluorescence intensities (0.1 sec integration time) were recorded following the incubation with 10 mM Ni^2+^, using constant excitation- and emission wavelengths of 492 and 512 nm, respectively, and 4 nm excitation- and emission slit widths. For ion titration experiments, LeuT^tmFRET^ was incubated in fluorescence buffer containing 750 µM Ni^2+^ and increasing concentrations of Na^+^, K^+^ or Rb^+^ (0-1584 mM), substituted with Ch^+^ to preserve ionic strength. Fluorescence intensities were obtained using a FluoroMax-4 spectrophotometer and the same technical specifications as described above. For Ni^2+^ titration experiments performed in 96-well plates (Corning), LeuT was subjected to increasing concentrations of Ni^2+^ (10^-7^ to 10^-2^ M) in buffer containing 800 mM of NMDG^+^, Na^+^ ± 50 µM leucine or K ± 50 µM leucine. Fluorescence intensities at 512 nm were recorded on a PolarStar Omega plate reader (NMG Labtech) upon excitation at 492 nm.

### Analysis of tmFRET Data

Fluorescence intensities for LeuT^tmFRET^ variants (F) were normalized to their equivalents for LeuT^K398C^ (F_no site_) without the metal-chelating His-X_3_-His site, to correct for the contribution of collisional quenching from free Ni^2+^, dilution, as well as the primary inner filter effect (1-F/F_no site_). An increase in 1-F/F_no site_ implies an enhanced energy transfer between FRET-donor (F5M) and -acceptor (Ni^2+^) that, when plotted as a function of increasing Ni^2+^, was fitted to a Hill equation from which EC_50_ and maximal tmFRET values were obtained. Maximal tmFRET efficiencies were converted to distances by the FRET equation:

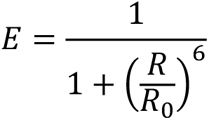

With *E*, *R* and *R_0_* being the efficiency of energy transfer, distance between FRET probes and Förster distance (*R_0_* = 12 Å for Ni^2+^/fluorescein^38^), respectively. For ion titration experiments, fluorescence intensities for LeuT^tmFRET^ (not normalized to LeuT^K398C^) at each ion concentration (F) were normalized to fluorescence intensities obtained in the absence (F_0_) of Na^+^, K^+^ or Rb^+^ (F_0_/F). As for 1 – F/F_no site_, F_0_/F is inversely dependent on the distance between the FRET probes and when plotted against the ion concentration, data points could be fitted to a Hill equation, yielding the Hill slope and EC_50_ value. Experiments were repeated at least three independent times in triplicates using protein from two separate purifications. Data points and Hill slopes are reported as mean ± s.e.m., whereas EC_50_ values are mean [s.e.m. interval]. When indicated, data points were subjected to statistical analysis using either *post hoc* Tukey or Bonferroni multiple comparison test as part of a one-way analysis of variance (ANOVA).

### Reconstitution of LeuT in liposomes

*E. coli* polar lipid extract dissolved in chloroform (Avanti Polar Lipids) was dried under a steam of N_2_ for 2 hours. Lipids were re-suspended to 10 mg/ml in reconstitution buffer (200 mM NMDG-Cl, 20 mM Tris/HEPES (pH 7.5)) by vortexing and 2x 10 min bath sonication. The lipid solution was subjected to 5 freeze-thaw cycles in ethanol dry ice bath. Subsequently, the liposomes were extruded with a mini extruder (Avanti Polar Lipids) 11 times through a Nuclepore^TM^ Track-Etch Membrane polycarbonate filter of pore size 400 nm (GE Healthcare Life Siences) and diluted to 4 mg/ml in reconstitution buffer. Liposome destabilization was induced by stepwise addition of 10 µl aliquots of 10 % (v/v) Triton X-100. Destabilization of liposomes was followed by measuring absorbance at 550 nm. Triton X-100 was added until absorbance of the sample had reached a maximum and started to decrease. LeuT solubilized in DDM was added in a protein to lipid ratio of 1:25 (w/w) and the protein-liposome solution was incubated under slow rotation for 30 min at 4℃. Semidry SM-2 Bio-Beads (Bio-Rad Laboratories), equilibrated in reconstitution buffer, were added in aliquots of 8.5 mg beads/mg lipid after 30 min, 60 min, 120 min and after overnight incubation at 4℃ with gentle agitation. The beads were filtered out 2 hours after the last addition of bio-beads. The proteoliposome solution was diluted ∼20 times in the indicated internal buffer (200 mM salt (NaCl, KCl, NMDG-Cl, CsCl or RbCl), 20 mM Tris/HEPES (pH 7.5)) and centrifuged for 1 h at 140,000 *g* at 4℃. Pelleted proteoliposomes were re-suspended in the indicated internal buffer to a final concentration of 10 mg lipid/ml. Single-use aliquots of proteoliposomes were flash frozen in liquid N_2_ and stored at -80℃ until further use.

### [^3^H]Alanine uptake into proteoliposomes

Proteoliposomes were thawed and extruded through a Nuclepore^TM^ Track-Etch Membrane polycarbonate filter with 400 nm pore size (GE Healthcare Life Siences). Uptake was assayed in a 96-well setup in ultra-low attachment, round bottom plate (Costar) at room temperature (21-23℃). Uptake buffer (20 mM Tris/HEPES (pH 7.5), 200-225 mM salt (NaCl, KCl or NMDG-Cl)) supplemented with [^3^H]alanine (Moravek Biochemicals) was added to each well in a volume of 190 µl. Subsequently, 10 µl of proteoliposome solution was added to start the uptake reaction. For time-dependent uptake, the uptake buffer was supplemented with 2 µM [^3^H]alanine with a specific activity of 1.335 Ci/mmol and proteoliposomes were added at the indicated time points (2 - 60 min). For concentration dependent uptake, uptake buffer was supplemented with [^3^H]alanine with a specific activity of 3.73 Ci/mmol (50 - 2000 nM) or 1.07 Ci/mmol (3.5 – 6 µM). The uptake reaction was terminated after 5 min. To assess nonspecific [^3^H]alanine uptake and binding, proteoliposomes were pre-incubated with 200 µM unlabeled leucine for 15 min, and 100 µM unlabeled leucine was added to the uptake buffer to saturate all transporters with leucine. The uptake reaction was terminated by filtering the samples through a 96 well glass fiber filter (Filtermat B – GF/B, Perkin Elmer) soaked in 1.5% poly(ethyleneimine) solution, using a cell harvester (Tomtec harvester 96 match II). The filter was washed with 1 ml ice-cold wash buffer (200 mM ChCl, 20 mM tris/HEPES (pH 7.5)) for each of the 96 positions on the filter. Filter plates were dried at 96℃ before scintillation sheet (MeltiLex B/HS, Perkin Elmer) was melted on to the filter. Filters were counted on a 2450 MicroBeta^2^ microplate counter (PerkinElmer) in “normal” counting mode.

### SDS-PAGE analysis of LeuT orientation

As for uptake experiments, proteoliposomes were extruded through a Nuclepore^TM^ Track-Etch Membrane polycarbonate filter with 400 nm pore size (GE Healthcare Life Siences). Reconstituted LeuT corresponding to 8 µg was treated with thrombin (0.5 unit; Cytiva) and the reaction was quenched upon the addition of 500 µM PMSF after 1, 5, 10, 30, 60, 120 and 180 min. Controls without thrombin and with 1.5 % DDM (for 180 min) were included. Following sodium-dodecyl-sulfate polyacrylamide gel electrophoresis (SDS-PAGE), the gel was stained with InstantBlue (abcam) and images were obtained using an ImageQuant 800 (Cystiva).

### Radioactive binding assay for proteoliposomes

To assess the amount of active protein in each reconstitution condition, a sample of proteoliposomes from each of the different intra-vesicular buffer conditions was solubilized in buffer (30% glycerol, 1 % (wt/vol) DDM, 20 mM Tris/HEPES (pH 7.5), 200 mM NaCl). The samples were left for 3 h with gentle agitation at 4℃ to dissolve proteoliposomes and solubilize LeuT in DDM detergent micelles. The amount of re-solubilized active LeuT was assessed by binding in a saturating [^3^H]leucine concentration (1 µM) in buffer (200 mM NaCl, 20 mM Tris (pH 8), 0.05 % (w/v) DDM) by SPA. Measures of maximum binding from each condition were used to normalize the c.p.m. obtained from uptake experiments to ensure that uptake was not affected by variations in protein content in the proteoliposome samples. The highest maximum binding was used as normalization standard.

**Analysis of [^3^H]alanine uptake data**. For **time-dependent uptake**, specific uptake data, normalized to the amount of active protein (see section above), was fitted to a one-phase association using GraphPad Prism 9.0.

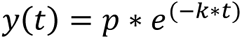

Where *p* is the maximal uptake (the amplitude) and *k* is the rate constant in min^-1^, *t* is time in min and *y* is uptake in c.p.m. after normalization. Data from each experiment was subsequently normalized to the value of *p* form the condition with NMDG^+^. The normalized data sets from each experiment were combined and re-fitted to a one-phase association.

For **concentration-dependent uptake**, normalized, specific uptake data was fitted to the Michaelis-Menten equation:

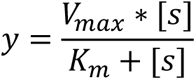

Where *V_max_* is the maximal uptake rate, *K_m_* is the substrate concentration where half-maximum uptake rate is reached, *y* is the uptake in c.p.m. after normalization. *[s]* is the [^3^H]alanine concentration in µM. Subsequently data was either combined or (if shown in % on the y-axsis) normalized to the *V_max_* value determined with intra-vesicular NMDG^+^. The normalized data sets from each experiment were combined and fitted to the Michaelis-Menten equation. All uptake experiments were done in triplicates or duplicates as indicated and repeated 3 - 4 times using protein obtained from at least 2 different purifications and reconstitutions.

For concentration dependent uptake with different intra-vesicular cations, c.p.m. were converted to d.p.m., and then to pmol of [^3^H]alanine per min., using the specific activity of [^3^H]alanine and a counting efficiency of 40% for 2450 MicroBeta^2^ microplate counter (PerkinElmer) in “normal” counting mode.

## ACKNOWLEDGEMENTS

We would like to thank Patricia Curran for technical guidance. We also acknowledge the NINDS intramural program for support.

## AUTHOR CONTRIBUTIONS

CJL conceptualized the ideas and designed the experiments together with SGS, AN and JAM. SGS and AN performed the experiments and did the data analysis together with CJL. All authors were involved in data interpretation. SGS, AN and CJL prepared the manuscript all authors read and commented on it.

## COMPETING INTERESTS

The authors declare no competing interests.

## FUNDING SOURCES

Support for this research was provided by the Independent Research Fund Denmark (1030-00036B to C.J.L.), the Lundbeck Foundation (R344-2020-1020 to C.J.L.), the Novo Nordic Foundation (NNF19OC0058496 to C.J.L.) and the Carlsberg Foundation (CF20-0345 to C.J.L.).

---

**Figure 1 – Figure supplement 1.**
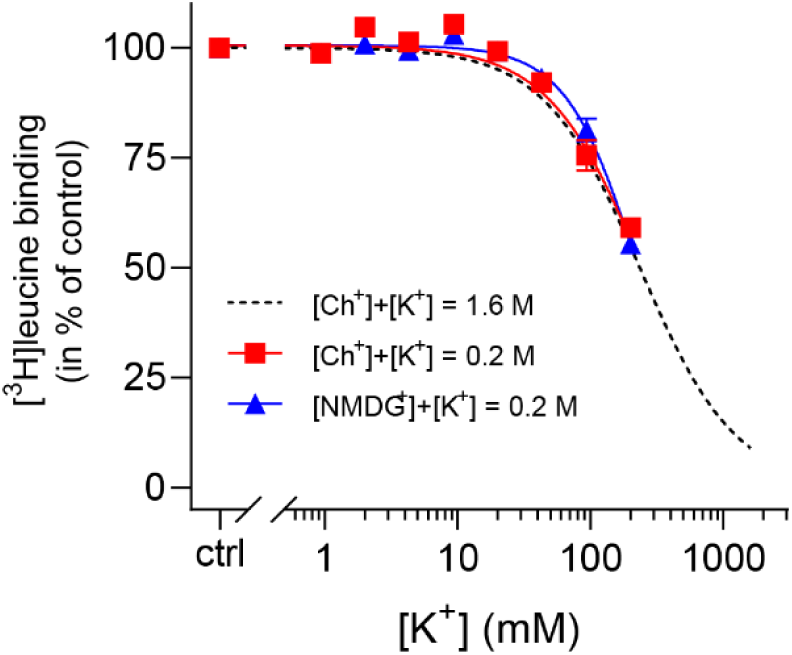
K^+^ inhibition is preserved for LeuT in low ionic strength. (**A**) Displacement of Na^+^-mediated [^3^H]leucine (10x *K*_d_) binding to LeuT by 0-200 mM K^+^, using Ch^+^ (red squares) or NMDG^+^ (blue triangles) as the counter ion. Displacement with 0-1600 mM K^+^, using Ch^+^ as counter ion, is shown for reference (black dashed line, from Figure 1B). The assay is performed in the presence of 7.7 mM Na^+^. Data are normalized to a control with 0 mM K^+^ and modelled by the Hill equation. Data are shown as mean ± s.e.m. (error bars often smaller than data points), n= 3–6 conducted in triplicates. The IC_50_ values obtained using up to 0.2 M Ch^+^ or NMDG^+^ as the counter ions were not significantly different from that obtained in a total ionic strength of 1.6 M (Dunnett’s multiple comparison test (one-way ANOVA), *p* = 0.84 and *p* = 0.99, respectively).

**Figure 2 - Figure supplement 1.**
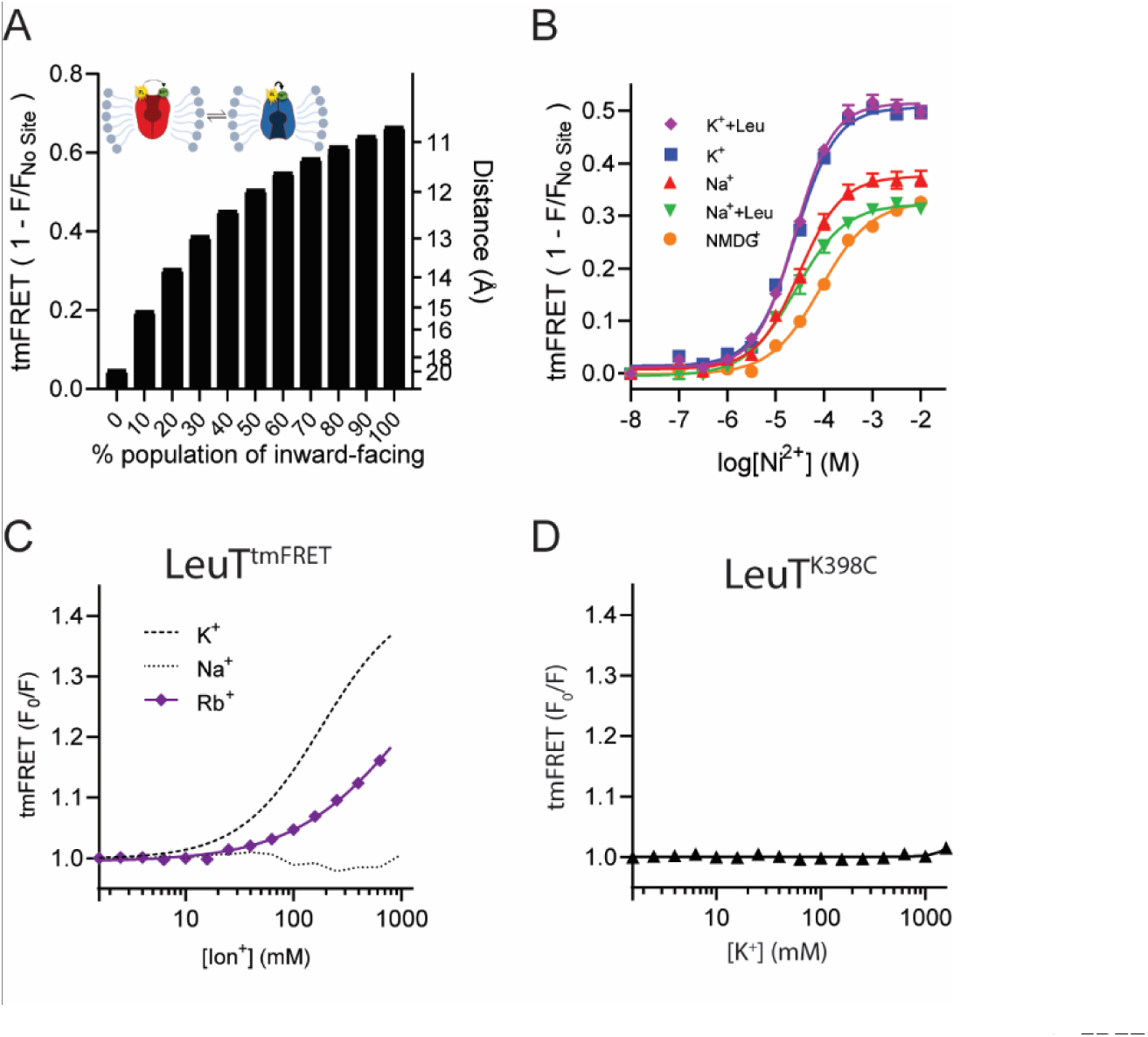
TmFRET principle and characterization of LeuT^tmFRET^. (**A**) Theoretical FRET efficiencies (1-F/F_no site_) (left axis), converted to distances by the FRET equation (right axis), as a function of the relative population of the outward closed state. This is according to R_obs_ = (p_OC_ R_OC_^-6^ + p_OO_ R_OO_^- 6^)^-6^, where p and R refer to the fractions and distances, respectively, for the outward-closed (OC) and outward-open (OO) states of LeuT^39^. The plot is based on the distances determined in (Figure 2A) between the sulfur atom of K398C (TM10) and the coordinated Ni^2+^ ion (EL4a) and relies on the assumption of a static distance distribution. (**B**) tmFRET efficiency (1-F/F_no site_) as a function increasing Ni^2+^ for LeuT^tmFRET^ incubated in 800 mM of K^+^ (blue squares, EC_50_ = 25.6 [24.8; 26.5] µM), K^+^ + 50 µM leucine (purple diamonds, EC_50_ = 24.6 [24.1; 25.1] µM), Na^+^ (red triangles, EC_50_ = 30.9 [29.1; 32.8] µM), Na^+^ + 50 µM leucine (green triangles, EC_50_ = 26.8 [23.1; 30.4] µM) and NMDG^+^ (black circles, EC_50_ = 85.1 [81.4; 88.8] µM). Data points are fitted to a Hill equation and EC50 values are given as mean [s.e.m. interval]. (**C**) tmFRET (F_0_/F) as a function of increasing Rb^+^ (purple) with Na^+^ (dotted line) and K^+^ (dashed line) for reference. Data points are fitted to a Hill equation. Data (A-C) are shown as means ± s.e.m., n = 3-4 in triplicates. (**D**) tmFRET (F_0_/F) as a function of increasing K^+^, performed on LeuT^K398C^ (lacking the His-X_3_-His site) in the presence of 750 µM Ni^2+^. Data points are shown as mean ± standard deviation (s.d.), n = 1 in triplicates. Error bars often smaller than data points.

**Figure 3 – Figure supplement 1.**
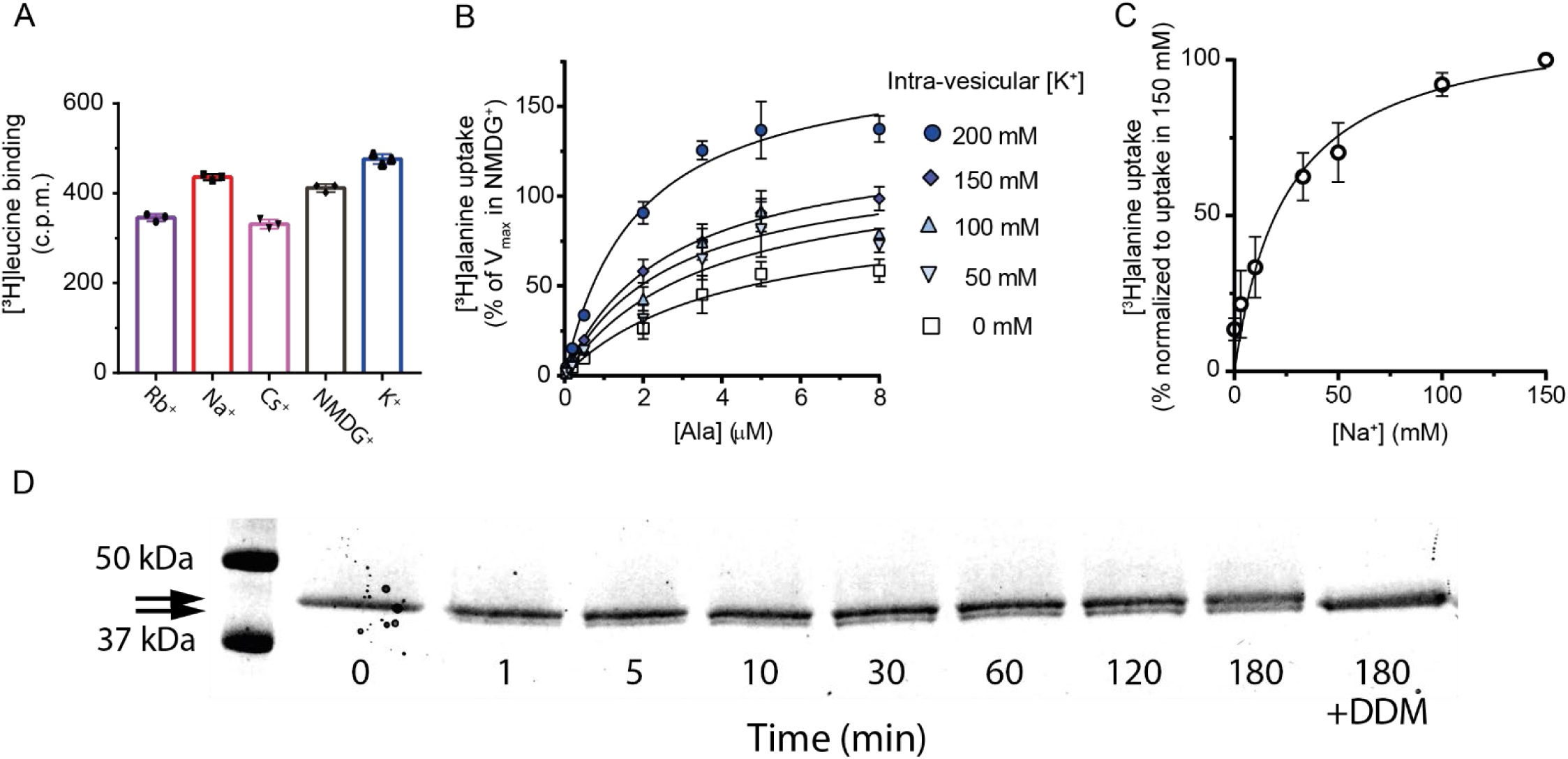
[^3^H]alanine uptake into proteoliposomes. (**A**) Binding of a saturating concentration (20x *K*_d_) of [^3^H]leucine to LeuT solubilized from the proteoliposomes. The amount of binding in counts per minute (c.p.m.) was taken as a measure of the relative number of active transporters in each reconstitution condition and used to adjust the uptake data in Figure 3B and C for relative reconstitution efficiency. Bar plots show the mean ± s.d. from a representative reconstitution performed in triplicates. (**B**) Concentration-dependent [^3^H]alanine uptake for 5 min in the presence of 200 mM Na^+^ into proteoliposomes containing increasing concentrations of K^+^. Lines are fits to Michaelis-Menten kinetics. The *V*_max_ in 0 mM K^+^ was defined as 100% (**supplementary table 3**). *V*_max_ increased with increasing intra-vesicular K^+^ (one-way ANOVA followed by a test for a linear trend, *p*=0.0001). None of the estimated *K*_m_ values were significantly different (Tukey multiple comparison-corrected one-way ANOVA). (**C**) [^3^H]alanine uptake for 5 min in the presences of varying Na^+^ concentration. Data are fitted to Michaelis-Menten kinetics. The Na^+^ concentration required to reach half-maximum uptake was 25 ± 8 mM (mean ± s.e.m.). Data in (B) and (C) are shown as mean ± s.e.m., n=3 in triplicates. **(D)** SDS-PAGE analysis of LeuT proteoliposomes following time-dependent thrombin digestion of accessible C-terminals (reducing the mass of LeuT by ∼1.3 kDa). The reaction was terminated by the addition of PMSF at the specified time points. The lanes corresponding to the time-dependent proteolysis are flanked by lanes containing proteoliposomes without thrombin (left, 0 min) or digested in the presence of DDM (right, 180 min+DDM). Arrows indicate bands of full-length (top) and cleaved (bottom) LeuT.

**Figure 3 - Supplementary table 1.**
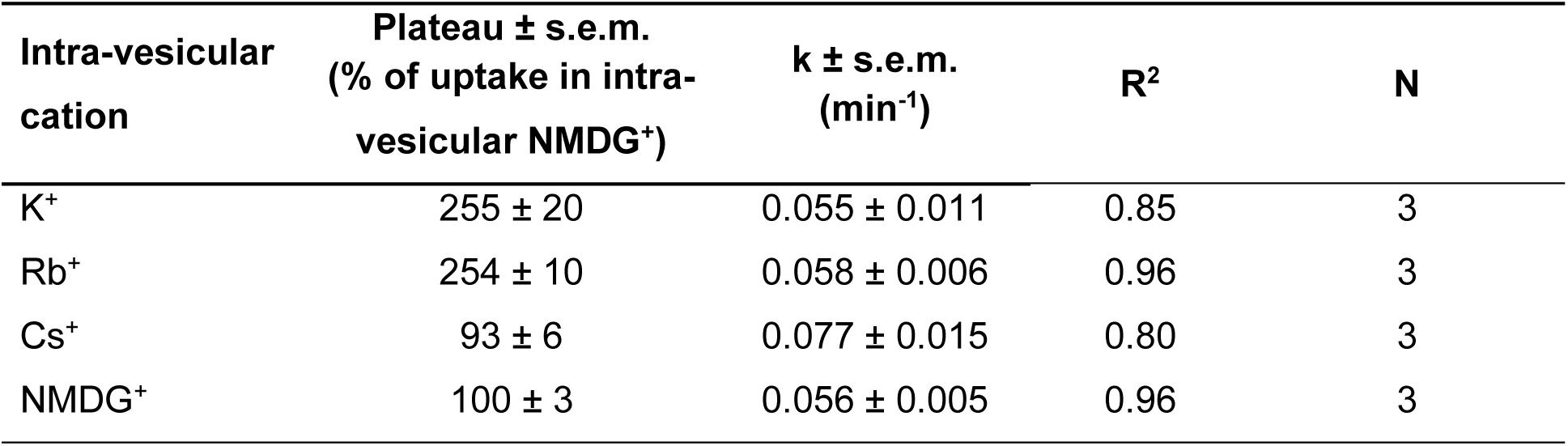
Rate constants for time-dependent uptake of [^3^H]alanine by LeuT into proteoliposomes. Rate constants from time-dependent [^3^H]alanine uptake into proteoliposomes with LeuT fitted to one phase association in GraphPad Prism 9.0 (see Figure 3B).

**Figure 3 - Supplementary table 2.**
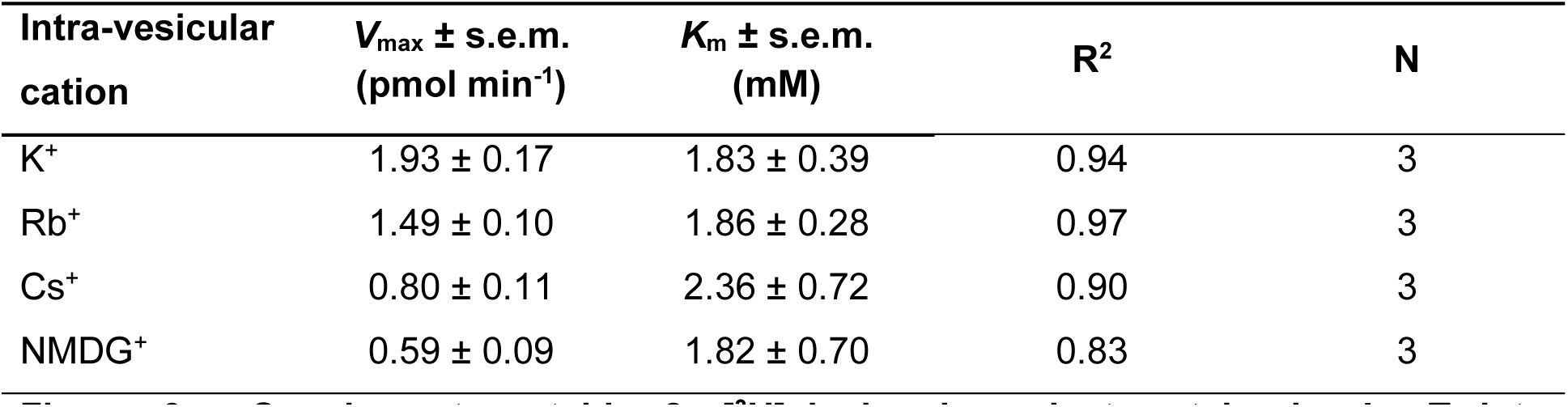
[^3^H]alanine-dependent uptake by LeuT into proteoliposomes. Constants from [^3^H]alanine-dependent uptake into proteoliposomes with LeuT containing increasing intra-vesicular [K^+^] fitted to Michaelis-Menten kinetics in GraphPad Prism 9.0 (see Figure 3C).

**Figure 3 - Supplementary table 3.**
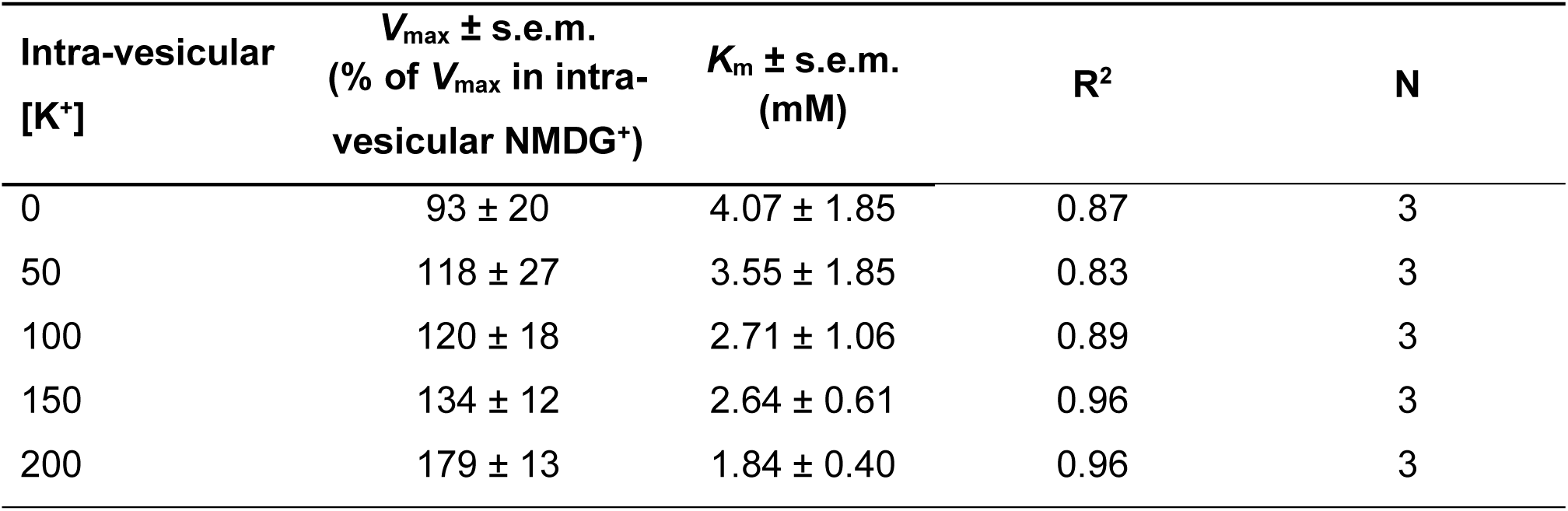
[^3^H]alanine uptake by LeuT into proteoliposomes with increasing intra-vesicular [K^+^]. Constants from [^3^H]alanine-dependent uptake into proteoliposomes with LeuT containing various intra-vesicular cations fitted to Michaelis-Menten kinetics in GraphPad Prism 9.0(see Figure 3 – Figure supplement 1B).

**Figure 5 – Figure supplement 1.**
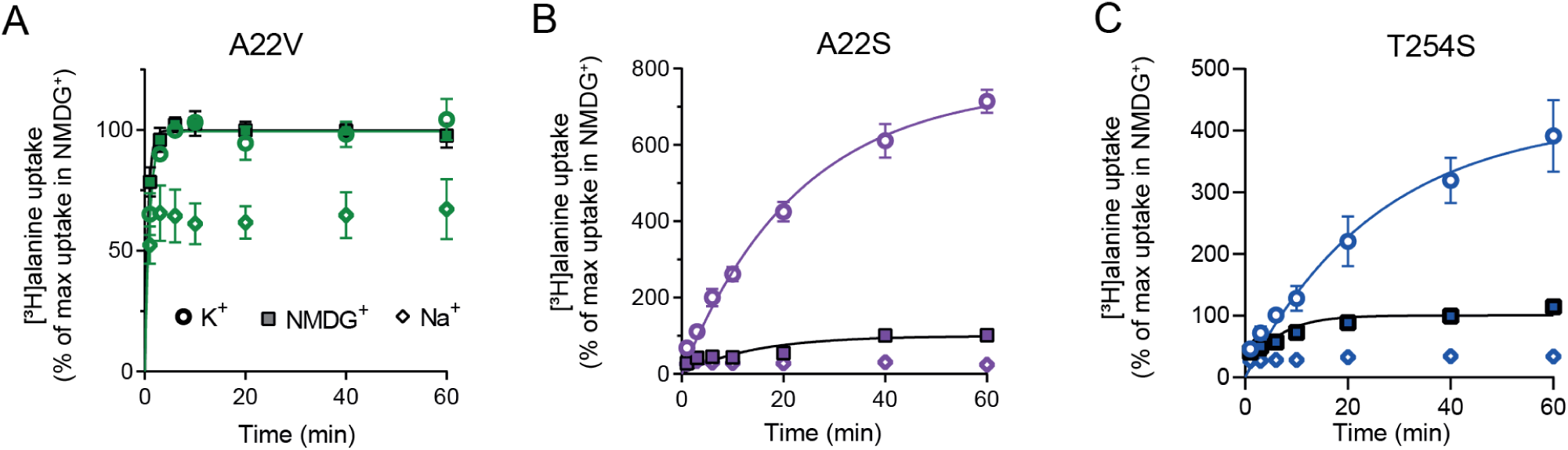
[^3^H]alanine uptake by selected Na1 mutants. (**A**) [^3^H]alanine activity over time by LeuT A22V in the presence of 200 mM Na^+^. LeuT A22V is reconstituted into proteoliposomes with either 200 mM Na^+^ (open diamond), NMDG^+^ (square) or K^+^ (open circle). Maximal transport obtained in NMDG^+^ was defined as 100%. Maximal [^3^H]alanine transport in K^+^ was 99 ± 2 %. The rate constant (k) was 1.55 ± 0.20 and 1.01 ± 0.15 min^-1^ in NMDG^+^ and K^+^, respectively, which correspond to a half-time to reach the concentrative capacity of 0.45 and 0.67 min. (**B**) [^3^H]alanine uptake by LeuT A22S. The [^3^H]alanine transport with K^+^ increased to 756 ± 37%. The rate constant (k) was 0.071 ± 0.02 and 0.044 ± 0.005 min^-1^ in NMDG^+^ and K^+^, respectively, corresponding to an half-time to reach the concentrative capacity of 9.8 and 15.8 min. **(C)** [^3^H]alanine uptake by LeuT T254S. Maximal uptake in K^+^ was 422 ± 51 %. The rate constant was 0.17 ± 0.04 and 0.04 ± 0.01 min^-1^ in NMDG^+^ and K^+^, respectively, corresponding to a half-time of 4.1 and 17.3 min. All data are shown as mean ± s.e.m. n = 3, performed in triplicates.

**Table 1 – Figure supplement 1.**
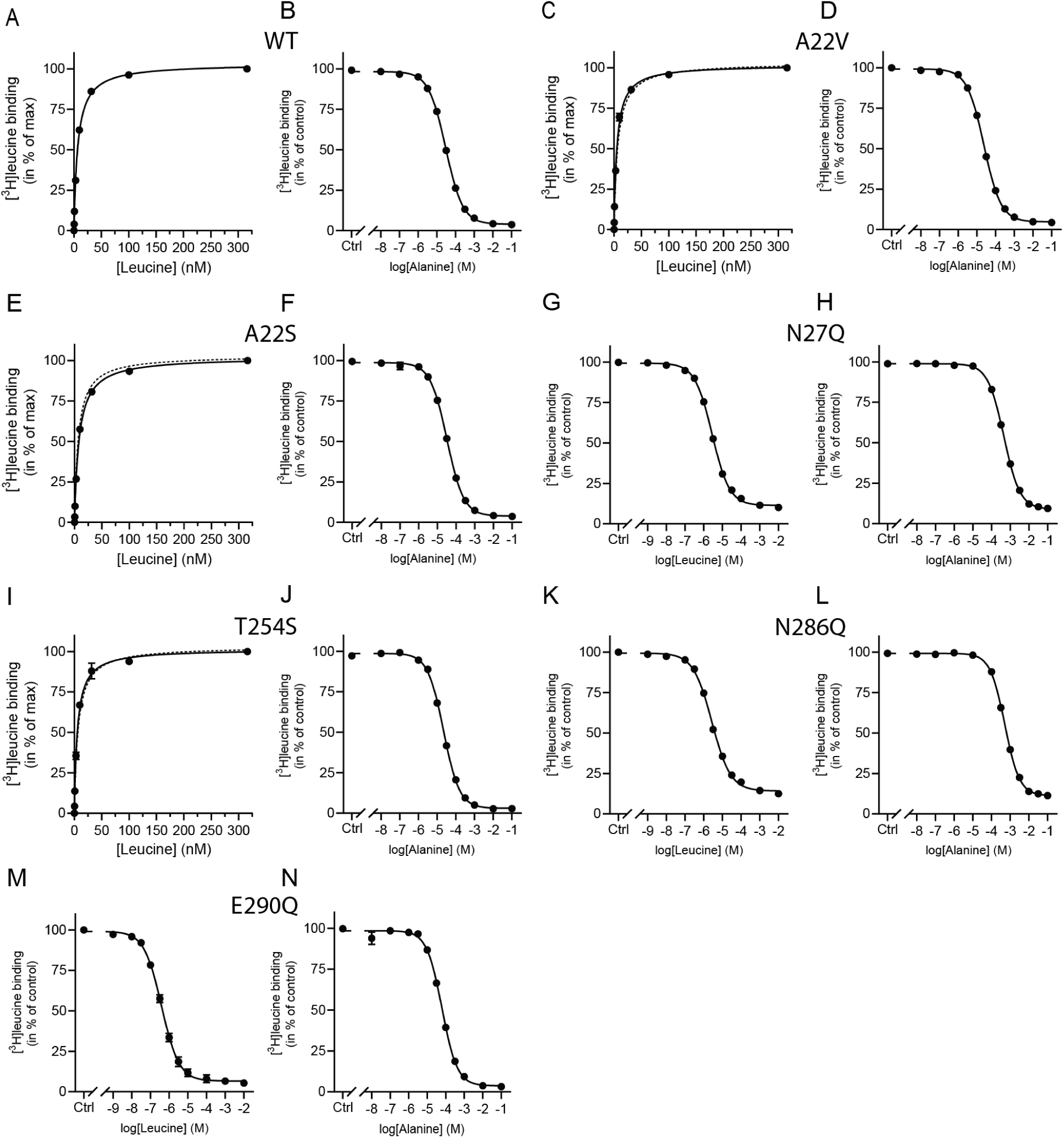
Leucine and alanine binding to Na1 site mutants. **(A**,**C**,**E**,**I)** [^3^H]leucine saturation binding for WT (A), A22V (C), A22S (E) and T254S (I). (**B**,**D**,**F-H,J-N**) Displacement of [^3^Hleucine by leucine for N27Q (G), N286Q (K) and E290Q (M), or by alanine for WT (B), A22V (D), A22S (F), N27Q (H), T254S (J), N286Q (L) and E290Q (N). Binding was assayed in 200 mM Na^+^. All data points are mean ± s.e.m., n = 3-4, and fitted to a one-site binding model (A, C, E, I) or non-linear regression fit. *K*_d_ and *K*_i_ values are summarized in Table 1.

**Table 2 – Figure supplement 1.**
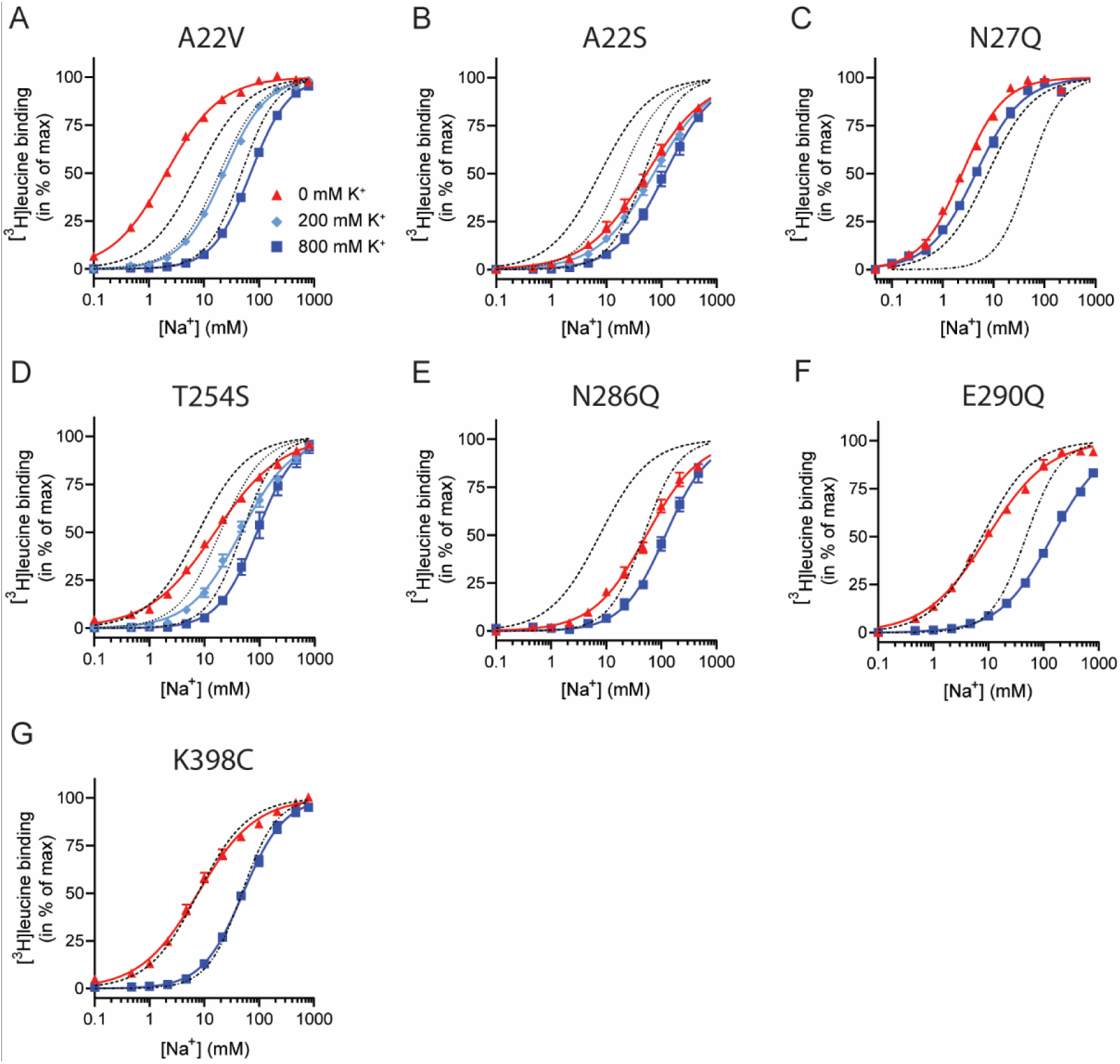
Na^+^-mediated [^3^H]leucine binding and inhibition by K^+^ by the LeuT mutants. (**A-F**) Na^+^-mediated [^3^H]leucine (10x *K*_d_) binding for the Na1 site mutants performed in the absence (red triangles) and presence of 200 mM (light blue diamond) and 800 (blue squares) mM K^+^. (**A**) A22V, (**B**) A22S, (**C**) N27Q, (**D**) T254S, (**E**) N286Q, (**F**) E290Q and (**G**) K398C. LeuT K398C was included as negative control. WT is shown for reference for 0 mM (dashed line), 200 mM (dotted line) and 800 mM (dash-dotted line) of K^+^. Ionic strengths were maintained using Ch^+^. All data points are shown as mean ± s.e.m. normalized to B_max_ and fitted to a Hill model. EC_50_ values are summarized in Table 2, n = 3-4.

## Notes

### Competing Interest Statement

The authors have declared no competing interest.

### Summary of Updates

Relative to the previous version, we have inserted a section that addresses the orientation of the LeuT molecules when reconstituted into the lipid bilayer membrane in the vesicles. The experiment showing the proportion of inside-out versus outside-out LeuT are shown in Figure 3- Figure Supplement 1D. We have also inserted a Figure 6, which illustrate a model for how K+ could influence the transport mode to avoid an exchange mode (efflux) and therefore increase the concentrative capacity and Vmax. We have included data for a non-Na1 site mutant in Table 2 and the experiment in Table 2- Figure supplement 1G. Also Figure 2- Figure Supplement 1B now has included tmFRET data on K++leucine as control. We have made other minor revisions and corrected a few typos.

